# How to Predict Effective Drug Combinations - Moving beyond Synergy Scores

**DOI:** 10.1101/2024.11.22.624812

**Authors:** Lea Eckhart, Kerstin Lenhof, Lutz Herrmann, Lisa-Marie Rolli, Hans-Peter Lenhof

**Affiliations:** Center for Bioinformatics, Saarland Informatics Campus, Saarland University, 66123 Saarbrücken, Germany; Computational Biology Group, Department of Biosystems Science and Engineering, ETH Zürich, 4056 Basel, Switzerland

## Abstract

To improve our understanding of multi-drug therapies, cancer cell line panels screened with drug combinations are frequently studied using machine learning (ML). ML models trained on such data typically focus on predicting synergy scores, which support drug development and repurposing efforts but have limitations when deriving personalized treatment recommendations. To simulate a more realistic personalized treatment scenario, we pioneer ML models that predict the relative growth inhibition (instead of synergy scores), and that can be applied to previously unseen cell lines. Our approach is highly flexible: it enables the reconstruction of dose-response curves and matrices, as well as various measures of drug sensitivity (and synergy) from model predictions, which can finally even be used to derive cell line-specific prioritizations of both mono- and combination therapies.

## Introduction

Tailoring drug treatments to the individual patient is a major goal of cancer research. Due to ethical concerns and limited availability of tumor material, relationships between molecular properties of cancer cells and their drug responses are generally not studied on humans directly, but instead using model systems, most prominently, cell lines. For monotherapy, large cell line panels such as the *Genomics of Drug Sensitivity in Cancer* (GDSC) database^1,2^ have been available for more than a decade, providing both molecular characterizations and drug screening data of cancer cell lines. However, combination therapies are frequently preferred over monotherapies for cancer treatment due to increased efficacy and a decreased risk of treatment resistance. ^3^ More recently, large data resources have also become available for drug combination screens: In 2019, the DrugComb data portal was introduced, ^4,5^ which accumulates harmonized results of drug screens from different sources. To date, a total of 37 datasets are available in DrugComb.^5^

Databases like the GDSC or DrugComb enable the systematic evaluation of the effect that different drugs have on different types of cancer cells. Thus, two main use cases that can be addressed using this data include (1) making personalized treatment recommendations for a given patient (cell line) and (2) finding promising drugs or drug combinations that should be further explored, e.g., for drug repurposing or the development of novel (combination) therapies. Due to the complexity and high dimensionality of the data, machine learning (ML) is commonly used to address these tasks.

ML models trained on monotherapy drug responses are usually suitable for both use cases, (1) and (2), since they directly predict measures of drug effectiveness, such as the IC50 or AUC value. In comparison, methods using drug combination data typically predict so-called drug synergy scores, ^6,7^ which are usually suited for the second task but less applicable for the first one as we briefly outline in the following: These scores quantify the synergistic or antagonistic potential of two compounds for a given cell line by comparing their combined effect on cell growth to the expected effect obtained from a baseline model that assumes no synergism or antagonism. ^8^ Prominent examples are the Loewe, ^9^ Bliss, ^10^ HSA, ^11^ and ZIP^8^ synergy score. For each of these scores, values *>* 0 indicate synergism, and values *<* 0 indicate antagonism, making it possible to classify drug combinations based on their synergy score. Undoubtedly, estimating the synergistic potential of compound combinations through synergy scores can be valuable for the identification of promising combination treatments to undergo more detailed screening, the development of novel compounds, or drug repurposing. However, even though synergy score prediction is sometimes motivated as a step toward achieving personalized treatment recommendations, ^6,12^ we believe that synergy scores have shortcomings that debilitate their usefulness for this application. Briefly summarized, synergy scores are based on various (in part very strong) model assumptions, some of which differ fundamentally between scores. ^8,13^ Additionally, disagreement between scores was observed by Vlot et al. ^13^ and Yadav et al., ^8^ weakening their informative value. Two factors that are especially relevant for personalized treatment recommendations are that (1) the scores are aggregated over multiple drug concentrations, which do not necessarily correspond well to clinically feasible concentration ranges^13^ (cf. Supplementary Figure 1) and (2) a high synergy between two compounds does not guarantee a high effectiveness of the combination treatment.^5^

Thus, instead of relying on synergy scores, we advocate exploring other strategies to estimate the effectiveness of combination treatments. For the prediction of drug combination sensitivity, several models that do not rely on synergy scores have been published: Malyutina et al. ^14^ and Zagidullin et al. ^4^ trained cell-line specific models that predict CSS (*Combination Sensitivity Score*) values, a sensitivity measure for two-drug combination therapies. ^14^ However, the CSS score is an aggregated measure of sensitivity based on drug-specific AUC values. Thus, like the AUC for monotherapies, ^15^ it depends strongly on the investigated concentration ranges and is not comparable across compounds.

Instead of focusing on one specific measure of drug sensitivity, an alternative approach is to directly predict the response (in terms of relative inhibition/viability) of cell lines at various treatment concentrations. Thereby, we could, moreover, reconstruct various drug sensitivity or synergy measures from the model predictions. For monotherapy, this approach has already been explored by Rahman and Pal et al. ^16,17^ For combination therapy, Zheng et al. ^5^ trained a CatBoost model that predicts the relative inhibition of two drugs at given concentrations for a given cell line. Similarly, comboFM by Julkunen et al. ^18^ employs higher-order factorization machines (HOFMs) to predict relative cell growth.

A drawback of all combination prediction approaches mentioned above is that they are not applicable to make predictions for previously unseen cell lines: Malyutina et al. ^14^ and Zagidullin et al. ^4^ trained cell line-specific models, while Zheng et al. ^5^ and Julkunen et al. ^18^ employ a one-hot encoding of cell lines and drugs in the model input such that both have to be known during training already. Thus, these models are difficult to apply for personalized treatment recommendations, where predictions should be made for a previously unseen patient (cell line). According to Codicè et al., this setting is frequently overlooked or insufficiently evaluated in ML-based drug response prediction. ^19^

In this manuscript, we present ML models for the prediction of drug combination sensitivity that do not rely on synergy scores and are able to make predictions for previously unseen cell lines, thereby mimicking a personalized treatment scenario. Instead of predicting an aggregated measure of treatment response, our models predict the relative inhibition at arbitrary treatment concentrations provided in the model input. Consequently, various measures of drug sensitivity or synergy, including dose-response curves and matrices, as well as IC50 values or synergy scores can be reconstructed from the model predictions.

We investigate not only different ML algorithms (neural networks, random forest, elastic net) but also analyze the benefit of including different drug characterizations (MACCS fingerprints, physico-chemical properties), as well as information on drug targets. The different model architectures provide different benefits, e.g., the ability to make predictions not just for two-drug combinations but also for monotherapies and combination treatments consisting of more than two drugs. Some of the investigated architectures also enable predictions to be made for any previously unseen drug, given that the features of the drug (e.g., MACCS fingerprint) are known. Our results show that random forests outperform the other algorithms in all investigated settings. Additionally, we analyze which inhibition intervals are predicted most accurately and investigate the reconstruction of mono- and combination sensitivity measures from our model predictions.

Lastly, using our recently published drug response measure called CMax viability,^15^ we showcase how our models can be applied to perform drug prioritization for mono- and combination therapies based on clinically feasible treatment concentrations. Drug prioritization, i.e., the ranking of drugs by their predicted effectiveness for a given cell line (patient) is a major goal in personalized medicine: it exceeds the mere prediction of sensitivity measures and moves toward deriving actual treatment recommendations.

## Materials and Data Processing

### Drug response data

Drug screening data for our analyses was obtained from the DrugComb database Version 1.5. More specifically, we employed the DrugComb API (https://api.drugcomb.org/) to download the list of all cell lines and their corresponding COSMIC IDs, the full list of drugs with their SMILE encodings and their target molecules, and the full dose-response matrices.

To assign the respective cell line and drug information to each dose-response matrix, we downloaded the core database from https://drugcomb.org/download, which provides a unique identifier for each dose-response experiment. Consequently, each database entry can be written as:

> (*cell line, drug row, drug col, conc row, conc col, inhibition*)

Here, *cell line* is the COSMIC ID of the investigated cell line, and *drug row* and *drug col* are the names of the tested drugs. The entries *conc row* and *conc col* are the micromolar concentrations of the tested compounds. For monotherapies, one of the drug names is set to *NULL* and the corresponding concentration is set to 0. Finally, *inhibition* denotes the relative inhibition measured after administration of the denoted drug concentration(s) (see Supplement for further information). Relative inhibitions *>* 0 denote reduced cell growth through the drug treatment, while inhibitions *<* 0 indicate increased growth.

We removed the following entries from the dataset:

- poor quality entries as defined by the authors of DrugComb^5^ with *inhibition <* −200 or *inhibition >* 200
- entries where the concentration of all tested drugs is 0 (*conc row* = *conc col* = 0)
- entries, where the corresponding cell line had no COSMIC ID or no gene expression data provided in the GDSC database

Additionally, we converted entries where *drug row* and *drug col* denote the same drug into monotherapies by summing the respective treatment concentrations and setting *drug col* to NULL:

> *(cell line, drug row, NULL, conc row + conc col, 0, inhibition).*

Cases where two different drugs are provided but only one has a concentration *>* 0 were modified to denote a monotherapy by replacing the drug with concentration 0 with *NULL*. Afterwards, all replicates involving the same cell line, the same drug(s), and same concentration(s) were averaged. Lastly, we log1p-normalized (*log*1*p*(*x*) = *log*(*x*+1)) the concentration values in *conc row* and *conc col*.

To keep the dataset size manageable, we only considered entries involving those 265 drugs (cf. Supplementary Table 1) for which at least 10,000 entries are provided after performing all the steps described above (cf. Discussion). Note that after this reduction still more than 10,000 entries remained for each of the drugs. In total, the final dataset consists of 5,291,424 entries covering 947 cell lines, 265 drugs, and 9,535 drug combinations.

Additionally, the CMax concentrations for 77 of the investigated drugs were obtained from Liston and Davis. ^20^ The CMax value denotes the peak plasma concentration after administering the highest clinically recommended dose of a drug. ^20^ In a recently published manuscript, we employed CMax to derive a novel drug sensitivity measure called the *CMax viability*, which will be described below. ^15^ We also use this measure to perform drug prioritization in the Results section.

### Drug Properties

For the representation of drugs in the inputs of our models, we investigated four different settings, which will be discussed below (cf. also Figure 1). Using the SMILES drug representations provided by DrugComb, we used RDKit version 2023.3.2^21^ to calculate two types of drug features:

- binary MACCS fingerprints^22^ of length 166
- 209 physico-chemical drug properties using the function CalcMolDescriptors from the rdkit.Chem.Descriptors module^23^

**Figure 1:**
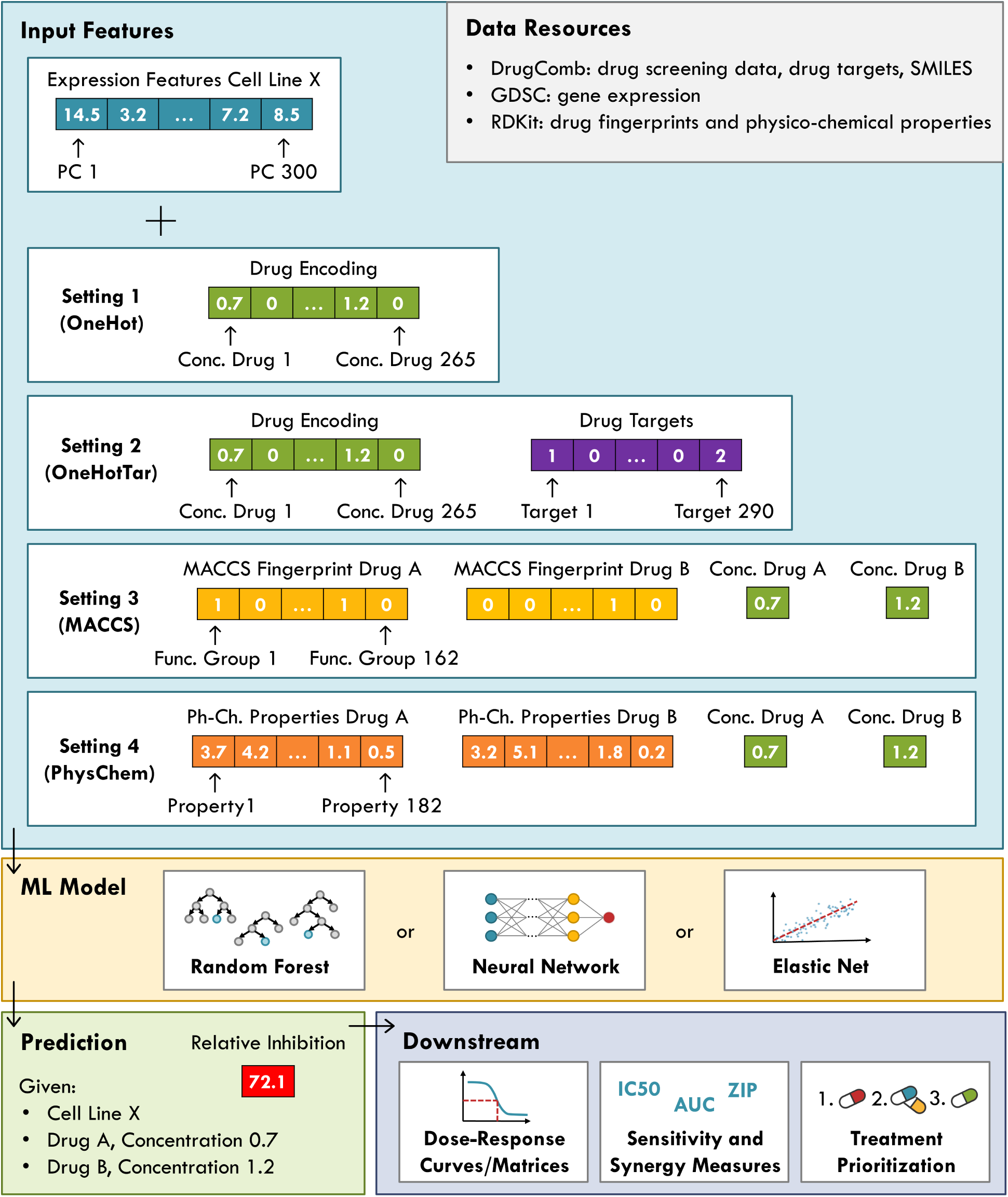
Prediction pipeline. This figure summarizes our pipeline for the prediction of relative inhibitions. The large blue box depicts the different types of input features and representations we investigated. The grey box at the top right lists our data resources. The yellow box shows the different ML algorithms we used. The green box at the bottom depicts the model output, i.e., the relative inhibition for a given cell-drug-drug combination at defined treatment concentrations. Lastly, the purple box shows potential downstream analyses that can be performed based on the model predictions.

We removed all properties that showed no variation across the investigated 265 drugs, resulting in MACCS fingerprints of length 162 and 182 physico-chemical properties.

Additionally, 735 drug target molecules for the investigated drugs were obtained from DrugComb.

### Gene Expression Data

Normalized gene expression data of 17,419 genes (*Affymetrix Human Genome U219 Array*) was obtained from the GDSC database Release 8.3 (https://www.cancerrxgene.org/downloads/bulk_download).

## Methods

### Model Inputs and Outputs

We train multi-drug models that predict the relative inhibition for a given cell line being treated with given concentrations of one or more drug(s). The model inputs comprise cell line features based on gene expression, a representation of the applied drugs, and the corresponding drug concentrations. For the representation of drugs, we investigated four different settings, which are depicted in Figure 1 and will be described below.

To characterize cell lines in the model input, we performed a principal component analysis (PCA) on the gene expression values of the training cell lines and used the first 300 principal components (PCs) as cell line features. This dimension reduction method performed well in our recently published benchmarking of drug sensitivity prediction methods. ^24^ The feature coefficients computed on the training data were used to project the test cell lines into the same 300-dimensional space. To perform the cross-validation discussed below, we re-computed the PCs based on the respective training folds.

In addition to the cell line features, we investigated four different settings for the encoding of drugs in the model input:

#### Setting 1 (OneHot)

In this setting, no drug properties are included. Instead, a 265-dimensional encoding of drugs is used. Each feature corresponds to one of the 265 drugs in our dataset. If a drug is part of the current entry, its feature is set to the corresponding log1p-normalized treatment concentration, otherwise it is set to 0.

#### Setting 2 (OneHotTar)

This setting uses the same concentration encoding as Setting 1 but additionally includes 290 drug target features. More precisely, we used the drug target annotations provided by DrugComb and included all molecules as targets that were targeted by at least five of the drugs in our dataset, resulting in a total of 290 target features. Each feature is then set to the number of drugs in the current entry that target the corresponding molecule (0, 1, or 2): Since DrugComb provides only data on monotherapies and two-drug combinations, the maximum value a target feature can have is 2, if it is targeted by both drugs in a two-drug combination entry. Note also that one drug can target more than one molecule.

#### Setting 3 (MACCS)

In this setting, each drug is represented by a 162-dimensional binary *molecular access system* (MACCS) fingerprint.^22^ Each position of the fingerprint corresponds to a molecular substructure, e.g., a functional group that may be present in a drug molecule. The respective bit is set to 1 if the corresponding substructure is present in the drug molecule at least once, and 0, otherwise. Additionally, one input feature for each drug is needed to denote its treatment concentration. Consequently, this setting uses a total of 2 · 162 + 2 · 1 = 326 drug features. To encode monotherapies, one of the fingerprints and the corresponding concentration are set to 0.

#### Setting 4 (PhysChem)

This setting is similar to Setting 3 but replaces each MACCS fingerprint with 182 numerical physico-chemical descriptors that denote different properties of the respective drugs, such as the molecular weight, number of valence electrons, or the logP value that measures lipophilicity. Consequently, this setting uses a total of 2 · 182 + 2 · 1 = 366 drug features. To denote monotherapies, one set of properties and the corresponding concentration are set to 0.

Depending on the desired application, the different settings provide different benefits: Settings 3 and 4 allow making predictions for arbitrary drug molecules given that their MACCS fingerprint or physico-chemical properties are known. Consequently, the resulting models can be used to make predictions for previously unseen, e.g., newly developed compounds. In contrast, models derived from Setting 1 and 2 are limited to those 265 drugs that were present in our dataset and hence encoded in the input. However, these models can not only make predictions for single drugs and two-drug combinations but even for treatments using three or more drugs simultaneously. While three-drug combination therapies have already been approved for cancer treatment by the United States Food and Drug Administration (FDA),^25^ DrugComb does not provide such data.

### Machine Learning Algorithms

We investigate the predictive performance of three ML algorithms: neural networks random forests, and elastic net. We chose these models, since neural networks and tree-based methods are commonly used for synergy prediction. ^7^ Furthermore, neural networks are also popular for drug sensitivity prediction, ^26–28^ while random forest and elastic nets are used less frequently for this task.^29–34^ In our recently published benchmarking, we found, however, that tree-based methods and elastic nets frequently outperform neural networks in predicting drug responses. ^24^ In line with our findings, several studies found that deep learning does not improve over conventional ML algorithms for making predictions on tabular data,^35–37^ or to generate feature representations for model inputs. ^24,38^

All prediction models were implemented in Python 3.11. Random forests and elastic net models were implemented using scikit-learn Version 1.5.0, ^39^ while neural networks were implemented using tensorflow Version 2.16.1^40^ with GPU support. The hyperparameters for each algorithm are provided in Table 1.

**Table 1:** Hyperparameters of the investigated ML algorithms. This table denotes the tuned hyperparameters for each ML algorithm. For hyperparameters not stated explicitly, the default parameters as provided the respective Python package were employed. Explicitly tuned hyperparameters are marked in bold. For the PhysChem setting (i.e., the setting with the largest data matrix), we were unable to train neural networks with the ELU activation or learning rates of 0.1 due to insufficient memory for resource allocation even when decreasing the batch size.

| Model | Parameter | Value(s) |
| --- | --- | --- |
| Elastic net | **alpha** | 0.01, 0.1, 1, 10, 100 |
|  | **l1_ratio** | 0, 0.25, 0.5, 0.75, 1 |
| Random forest | **max_depth** | 100, 1000000 |
|  | **max_features** | 25, 50, 100, 250 |
|  | **min_samples_leaf** | 2, 20, 100, 1000 |
|  | n_estimators | 500 |
| Neural network | loss | mean_squared_error |
|  | **activation** | tanh, ELU (none in last layer) |
|  | optimizer | Adam |
|  | **learning_rate** | 0.0001, 0.001, 0.1 |
|  | **hidden_layers** | 1,2,3,4,5 |
|  | size of hidden layers | equally spaced btw. in-/output size |
|  | **dropout** | 0.1, 0.3 |
|  | batch_size | 256 |
|  | bias_initializer | 0.01 |
|  | kernel_initializer | glorot_uniform for tanh,<br>he_normal for ELU activation |
|  | kernel_regularizer | l2 |
|  | epochs | 300 |
|  | validation_split | 0.2 |
|  | early stopping | yes |
|  | patience | 15 |
|  | restore_best_weights | True |

### Model Training and Testing

After filtering and processing the data as described above, we randomly divided the remaining cell lines into a training set (80% of cell lines) and a test set (20%). Since multiple data entries exist for each cell line (screening of different drugs/drug combinations at different concentrations), the final training data consists of all entries involving a cell line from the training set (3,741,209 entries). The final test data contains all remaining entries (1,550,215), i.e., all entries involving a cell line from the test set. This splitting ensures that the test performance is always evaluated on cell lines that were unseen during model training, thereby mimicking the scenario of making predictions for a previously unseen patient. In contrast, the same drugs and drug-combinations can occur in both the training and test data.

On the training data, we performed a 5-fold cross validation (CV) to determine the best-performing hyperparameters of each ML model (see Table 1). The CV folds were generated by randomly dividing the training cell lines into five disjoint folds and assigning all entries involving a certain cell line to the corresponding fold. Since the number of available entries per cell line differs, the size of CV folds varies slightly between 644,308 and 857,361 entries. For the hyperparameter combination with smallest mean absolute error (MAE) averaged across all five folds, one final model is trained on the complete training data and its performance is evaluated on the test data.

For the models using one-hot encodings (Setting 1 and Setting 2), each drug has a designated input node. This is not the case for the models using drug features (Setting 3 and Setting 4), where swapping the features and concentration of the first drug with those of the second drug represents the same treatment but results in changes in the input representation (cf. input visualization in Figure 1). However, the model output should not depend on the order of the drugs in the input, i.e., it should not depend on whether drug features of a drug A in the input vector are located in front of or behind those of a drug B. Therefore, each original sample is included twice in the datasets for Settings 3 and 4. These duplicate samples differ only in the order of the drug features and concentrations: once in the order A-B, once in the order B-A. In the Results section, we investigate the impact on model performance when models are trained using the duplicated versus non-duplicated data. The test performance is always evaluated on the duplicated entries.

### Fitting of Dose-Response Curves and Computation of Sensitivity Measures

Using the relative inhibitions predicted by our models, it is possible to reconstruct doseresponse curves for monotherapies and dose-response matrices for combination therapies (cf. Figure 2). Based on these curves/matrices, various measures of drug response can be derived. To this end, we first converted the (actual and predicted) relative inhibitions into relative viabilities by subtracting the relative inhibitions from 100 and dividing the result by 100. Additionally, we clamped viabilities to [0, 1]. Note that we report relative viabilities in range [0, 1] rather than range [0, 100] to keep the results consistent and comparable to our previous study. ^15^

**Figure 2:**
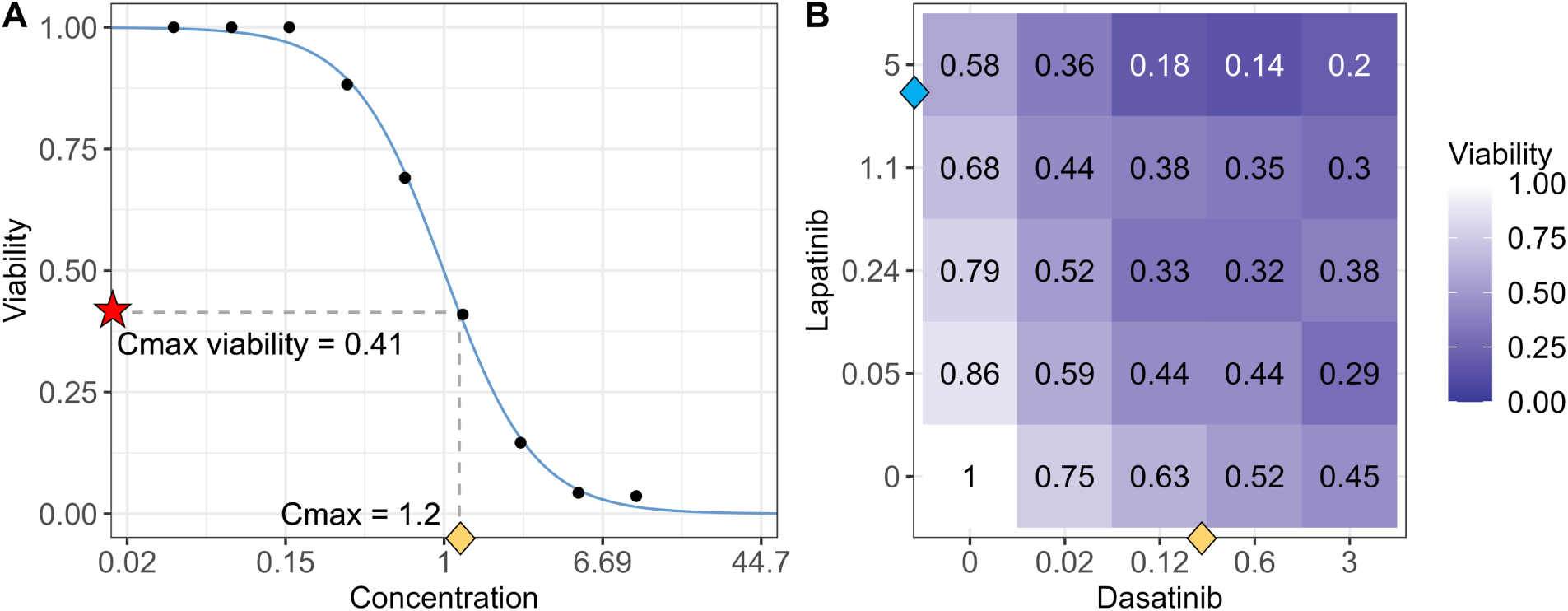
Exemplary dose-response curve and matrix. Sub-figure A depicts a dose-response curve (blue) for the monotherapy treatment of a cancer cell line (COSMIC ID 683667) with the drug Vorinostat. The fit is based on nine dose-response points (black). The yellow diamond marks the CMax concentration of Vorinostat (1.2*µ*M), and the red star marks the corresponding CMax viability (0.41) derived from the curve (cf. Methods). Sub-figure B depicts a dose-response matrix for the combination treatment of cell line 909755 with Dasatinib and Lapatinib, where the x- and y-axes denote the respective treatment concentrations. The yellow and blue diamonds approximately mark the CMax concentration of both drugs, which are used to limit the considered concentration combinations for the computation of the combination CMax viability (cf. Methods).

To perform the curve-fitting for monotherapies, we employed a three-parametric logistic function from the drc R-package: ^41,42^

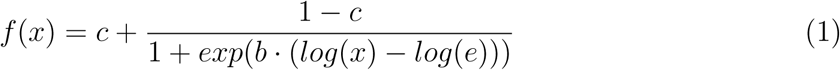

Here, *f* (*x*) denotes the estimated relative viability of the considered cell line at drug concentration *x*, *c* denotes the curve asymptote for increasing concentrations, *b* denotes the curve’s slope, and *e* denotes the concentration at the inflection point. We only fit curves when at least five dose-response points were available and we discarded all curves where the root mean squared error (RMSE) between the actual viabilities and those derived from the curve was greater than 0.3, a threshold that was previously employed for the data generation in the GDSC database.^43,44^ From the fitted curves, we derived two measures of monotherapy drug responses, namely IC50 values and CMax viabilities. The CMax viability is a novel drug sensitivity measure which we recently published. ^15^ It is defined as the relative viability at the CMax concentration of the respective drug. The CMax concentration denotes the peak plasma concentration of a drug after administering the highest clinically recommended dose. ^20^ Thus, the CMax viability is designed to estimate the maximal effect a treatment can realistically achieve. For the computation of CMax viabilities, we evaluated the function of the fitted curve at the drug’s CMax concentration (cf. Figure 2A). For the computation of IC50 values, we intersected the dose-response curves with a horizontal line with y-intercept 0.5.

For combination therapies, we developed a variation of the CMax viability we call the *combination CMax viability* that can be derived from an actual/predicted dose-response matrix (cf. Figure 2B). Our initial idea was to interpolate the values in the dose-response matrix to derive the relative viability when administering the CMax concentration of both combination drugs simultaneously. However, two synergistic drugs may have certain concentration windows with particularly high synergy/effectiveness. ^45^ Thus, it is possible that the smallest viability is reached at a concentration combination smaller than the CMax concentrations. (Note that this should not happen for the dose-response curves we employed to compute the CMax viability for monotherapies since these curves are monotonically decreasing.) Consequently, we considered the entire concentration range below the respective CMax values to compute our sensitivity measure. Conceptually, we want to derive the smallest viability within the area defined by the two concentration windows of the drugs limited at their respective CMax concentration. To compute the combination CMax viability, we linearly divided the concentration interval from 0 to the CMax for each drug into 100 equally spaced concentrations, each, resulting in 10,000 concentration combinations. For each combination, we estimated its relative viability through bilinear interpolation (R package pracma^46^) from the full dose-response matrix. Finally, we define the minimum of all 10,000 values as the combination CMax viability.

As the CMax denotes the maximal feasible treatment concentration for a drug monotherapy, it may not be feasible to administer the CMax concentration of two drugs in combination. Yet, we believe that the respective CMax concentrations are a reasonable upper bound to consider for the computation of combination CMax viabilities. Note also that administering the CMax concentration for monotherapies might likewise not be feasible in all cases. Furthermore, the presented approach can theoretically be applied to any desired concentrations other than CMax.

## Results

### Challenges of Using Synergy Scores for Personalized Treatment Recommendations

The idea behind synergy scores is to measure the synergistic or antagonistic potential of two compounds for a given cell line by comparing their experimentally measured combined effect on cell survival to the expected effect obtained from a baseline model that assumes no synergism or antagonism. ^8^ The baseline model is derived from monotherapy data of both compounds. It estimates their combined effect at the concentrations that were tested in the actual combination screening. The baseline and actually measured treatment responses are then subtracted from each other and the result is averaged over all concentration combinations to obtain a final synergy score. ^13^ Prominent examples of synergy scores that differ solely in their computation of the baseline are the Loewe, ^9^ Bliss, ^10^ HSA, ^11^ and ZIP^8^ scores. For each of these scores, values *>* 0 indicate synergism and values *<* 0 indicate antagonism. A detailed description of the scores can be found in the Supplement.

Undoubtedly, estimating the synergistic potential of compound combinations through synergy scores can be valuable for the identification of promising combination treatments to undergo more detailed screening, the development of novel compounds, or drug repurposing. However, there are known limitations of synergy scores, which have been summarized and extensively discussed in a review by Vlot et al., ^13^ where they also performed several analyses using a large-scale drug combination dataset. Their findings can be briefly summarized as follows: Firstly, each synergy score is based on certain model assumptions, some of which might frequently be violated by real word data.^47,48^ For example, both the Loewe and ZIP score require fitting dose-response curves of a certain shape to the monotherapy data. The Loewe score furthermore requires both drugs to have the same minimum and maximum effect as well as a constant potency ratio. ^13^ In comparison, the Bliss score relies on the assumption that the combined effect of two non-interacting drugs is statistically independent. Even though pharmacological independence is not necessarily required to achieve statistical independence, ^13^ it is most likely that statistical independence is caused by pharmacological independence. However, due to crosstalk between biological processes affected by either drug, achieving true pharmacological independence may be unlikely.^48^

These examples also highlight that the assumptions between scores differ fundamentally. In their data analysis, Vlot et al. observed only a moderate to low correlation between the four different scores calculated on the same data, which might be explained by the different model assumptions. They also found that value ranges between scores are not comparable: the HSA and ZIP scores generally result in higher values than Loewe and Bliss. Additionally, Vlot et al. observed that synergy scores are relatively difficult to reproduce between replicated experiments, even though the measured drug responses used to derive the scores correlated well between replicates. Furthermore, while misclassifications (synergism vs. antagonism) between scores were rare, several scenarios were identified where scores are likely to disagree, which could typically be retraced to a violation of model assumptions.

Based on these findings, Vlot et al. advocate against the automated analysis of large-scale data using individual synergy scores. Instead, they recommend a careful investigation of the respective dose-response curves to then select an appropriate score. When training models that only predict synergy scores (instead of concentration-specific inhibitions/viabilities), this is hardly possible since we are unable to assess the underlying dose-response relationship to validate model assumptions.

We agree with these conclusions by Vlot et al. but would like to emphasize further points that make synergy scores difficult to use and interpret, especially for personalized treatment recommendations: A methodological criticism of synergy scores is that they are an aggregated measure over concentration ranges. The choice of meaningful concentration ranges is especially challenging for experimental drugs but crucial to draw meaningful conclusions for personalized medicine. We have previously shown that the screened concentration ranges in the GDSC database do not correspond well to clinically feasible treatment concentrations^24^ and similar observations can also be made for the DrugComb database (cf. Supplementary Figure 1). Another major factor that hampers the use of synergy scores for treatment recommendation is that a high synergy between two compounds solely implies that the combination treatment is more effective than the respective monotherapies. However, it does not guarantee an overall high effectiveness (in terms of large relative inhibition) of the combination treatment.^5^ It follows that synergy scores alone should not be used to compare the suitability of different treatment options for a given patient (cell line). In particular, synergy scores cannot be used to compare the effectiveness of different combination treatments. Furthermore, it is not possible to compare the effectiveness of combination therapies to monotherapies involving different drugs.

Based on these drawbacks of synergy scores in general and for treatment recommendation in particular, our analyses presented in the following focus on sensitivity prediction instead. Compared to the number of synergy prediction methods, ^6,7^ sensitivity prediction of drug combinations is understudied, especially when the goal is to make predictions for previously unseen cell lines as we have outlined in the Introduction section.

In the following, we analyze how accurately drug responses (here: relative inhibitions) can be predicted for combination therapies. We compare different ML algorithms and model inputs and investigate the reconstruction of sensitivity measures from the model predictions. Additionally, we show how both mono- and combination therapies can be ranked by their effectiveness for a given cell line using our recently developed sensitivity measure: the *CMax viability*.^15^

### Overall Performance Comparison

Figure 3 shows the performance of all investigated models in terms of test MAE (mean absolute error). The optimized hyperparameters for each model are provided in Supplementary Table 2. The first row depicts the results for the entire test data, while the second and third row focus on the data subsets representing mono- and combination therapies, respectively. Across all four settings, random forests resulted in the lowest error, followed by neural networks, while elastic net had the worst performance. An exception is the PhysChem setting, where neural networks were outperformed by elastic net.

**Figure 3:**
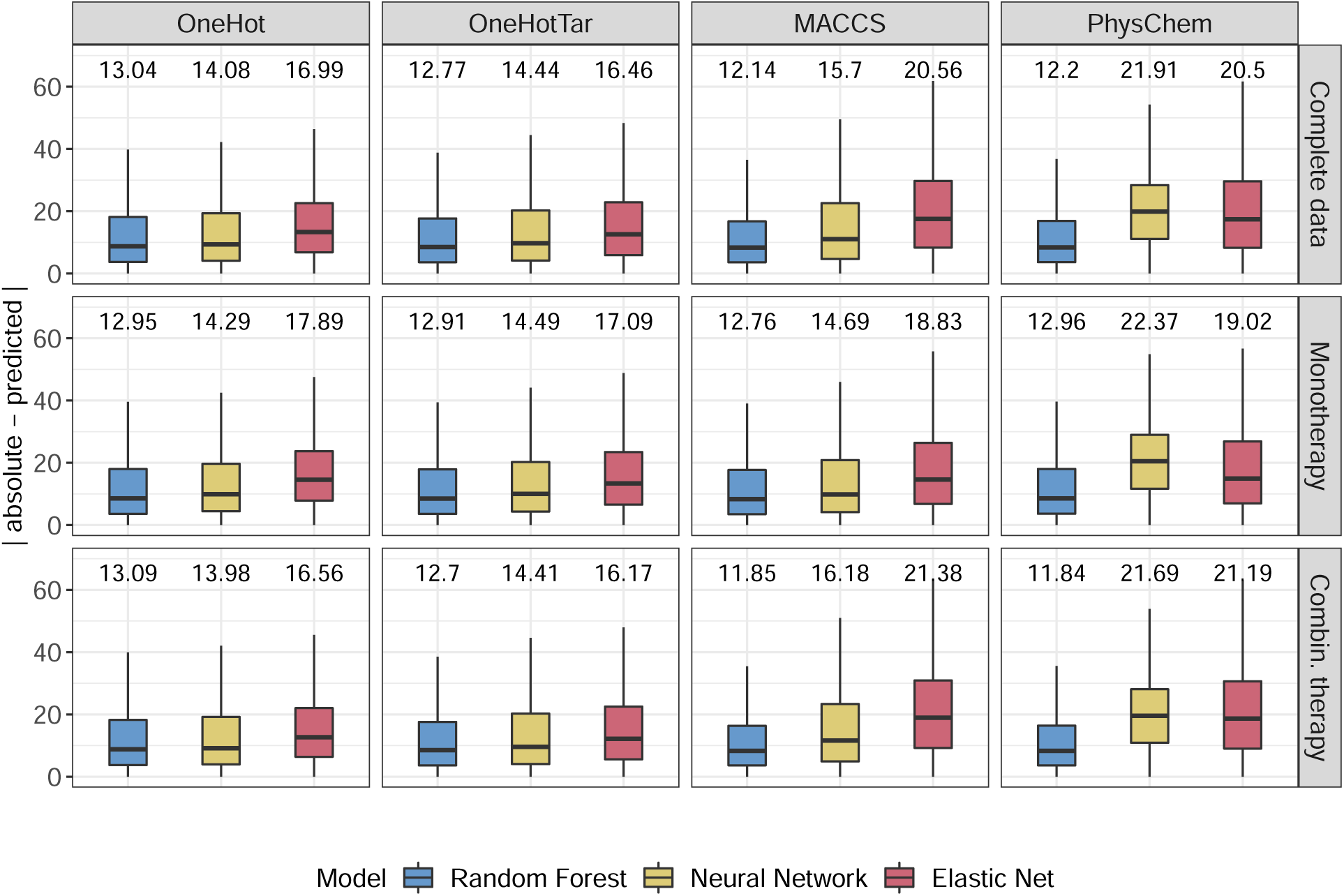
Test set performance. This figure shows the prediction errors (in terms of absolute difference between actual and predicted values) for each setting (columns) and each investigated ML algorithm (coloring). The first row shows the results for the entire test dataset, while the second and third row show the results for the data subsets corresponding to mono- and combination therapies, respectively. On top of each boxplot, the mean absolute error (MAE) is shown.

The overall smallest test error (MAE 12.14) was achieved using a random forest with MACCS fingerprints as input. Additionally, even the worst performing random forest model (OneHot, MAE of 13.04) still outperforms the best neural network (OneHot, MAE 14.08) and elastic net (OneHotTar, MAE of 16.46) models. Thus, the choice of ML algorithm seems to have a stronger impact on performance than the choice of input features, even though the different input representations differ considerably (cf. Methods and Figure 1). Notably, the addition of drug targets slightly improves predictions for random forest and elastic net but has the opposite effect for neural networks.

To contextualize the obtained errors, we compare them to two baseline models: A simple baseline model that always predicts the mean of the training data has a test MAE of 24.2. A more advanced baseline that always predicts the mean inhibition per drug for monotherapies and the mean inhibition of the combination for combination therapies has a test MAE of 19.74. Consequently, our best model improves these baselines by 50% and 37%, respectively. While all of the random forests models outperform the baseline, some elastic nets and neural networks are not superior to the baselines.

When investigating mono- and combination therapies separately (cf. row 2 and 3 of Figure 3), the same overall trends can be observed, with the random forest model with MACCS features again having the smallest error. Generally, both types of therapies can be predicted similarly well, even though the training data contains slightly more combination (60%) than monotherapy data (40%).

Besides the MAE, we also investigated the Pearson correlation (PCC) between the actual and predicted inhibitions. The overall PCC for the best-performing model was 0.8 (0.77 and 0.82 for mono- and combination therapies, respectively). However, computing correlations across the entire data artificially increases the PCC: since some drugs/combinations generally have lower/higher inhibitions than others, even mean predictions for each drug/combination (requiring no ML at all) would result in a correlation above 0. ^19^ Thus, we computed the mean per-drug PCC for monotherapies (0.58) and the mean per-combination PCC for combination therapies (0.56) (see also Supplementary Figure 2). These values have a similar magnitude to what we previously observed for monotherapy sensitivity prediction. ^15^

Note that Zheng et al. ^5^ and Julkunen et al. ^18^ also provide overall correlations and errors for the prediction of relative inhibition/growth (cf. Introduction). However, their results are not comparable to ours since we investigate the performance for unknown cell lines, which cannot be evaluated using the other two methods. It is known that the cell-line blind scenario increases errors considerably compared to making predictions for known cell lines. ^49,50^ To, nevertheless, assess how our random forest MACCS model would perform for known cell lines, we retrained the model using a random split of of the available data into a training (80%) and test set (20%). This split does not guarantee that cell lines in the test set were unseen during model training. Note that we still assured that duplicated entries denoting the same treatment are either exclusively contained in the training or the test set (cf. Methods). With a PCC of 0.96 and RMSE of 8.41, our performance for known cell lines is comparable to that reported by Zheng et al. (PCC = 0.98, RMSE = 7.12)^5^ and Julkunen et al. (PCC = 0.97, RMSE = 9.86 in cross-validation; PCC = 0.92 on validation data).^18^ However, the dataset used in our analyses is much larger and more heterogeneous comprising 947 cell lines, 265 drugs, and 9,535 drug combinations from different sources. In contrast, Zheng et al. employed solely the O’Neil dataset (39 cell lines, 38 drugs, 583 drug combinations), ^51^ which is known to be of high quality,^4,5^ whereas Julkunen et al. employed solely the AstraZeneca DREAM dataset (85 cell lines, 118 drugs, 910 drug combinations). ^6^

### Range Performance Comparison

Next, we investigated whether certain inhibition ranges can be predicted more accurately than others. Figure 4 shows the distribution of test MAEs for different inhibition intervals in range (−25, 100]. This range covers 99% of the training and test data. Predictions are (on average) most accurate in the interval (0, 25] followed by the interval (−25, 0]. As the actual inhibition increases, the error increases as well. This could be explained by the amount of available training data for each interval: Most data is located in the intervals (0, 25] (41%) and (−25, 0] (25%), while each of the other intervals is only covered by around 10% of the data. In Supplementary Section 3 and Supplementary Figure 3, we provide further analysis on how the amount of training data for individual drugs/combinations affects prediction performance.

**Figure 4:**
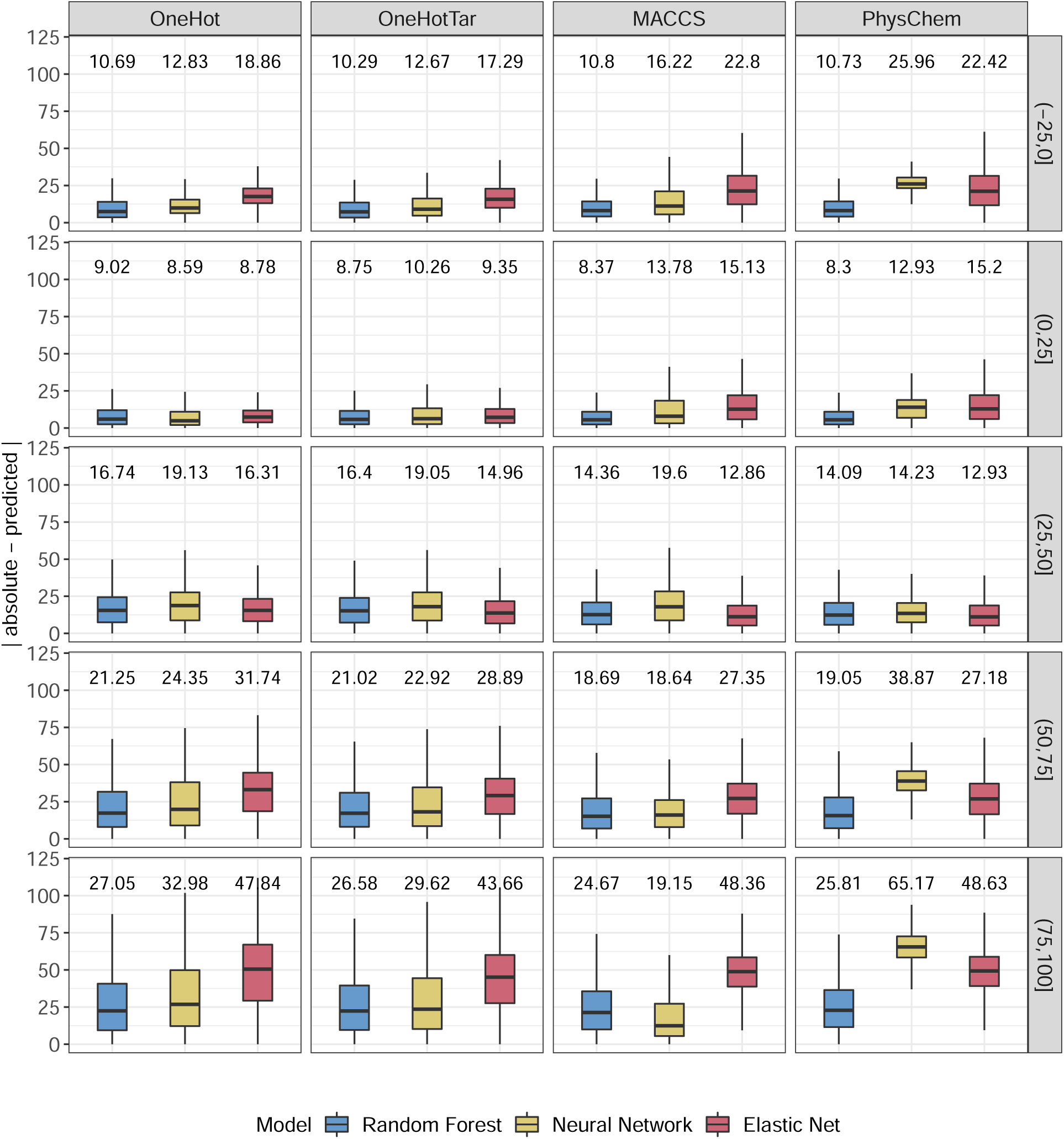
Test set performance for different inhibition ranges. This figure shows the prediction errors (in terms of absolute difference between actual and predicted values) for each setting (columns) and each investigated ML algorithm (coloring). Each row shows the performance for a different interval of actual relative inhibitions. On top of each boxplot, the mean absolute error (MAE) is shown.

Data points with high inhibition represent cases where the drug treatment greatly reduced the amount of viable cells, i.e., cases of effective treatment. Such data are commonly under-represented in drug screening datasets.^34,52,53^ They are, however, of particular interest for personalized therapy, where the most effective treatment options for a given patient should be determined.

Thus, for monotherapies, we developed SAURON-RF, a random forest-based model that is designed to improve predictions of drug-sensitive samples for both classification and regression. ^15,29^ To this end, SAURON-RF relies (among other things) on sample-specific weights. Consequently, we also tried to incorporate sample weights into our models presented here to increase the importance of the underrepresented intervals. Unfortunately, the sample weights had only little impact on predictions, especially for the cases with highest inhibition (see Supplementary Figure 4).

### Correlation of Duplicated Entries

As discussed in the Methods section, for the MACCS and PhysChem settings, the same treatment can be described by two different input representations through switching the order of the considered drugs (cf. Figure 1). Hence, we decided to include both input representations into the training and test data of our models. Ideally, predictions for both input representations should correlate well. Figure 5A shows the correlation of predictions for the best-performing random forest model trained using MACCS fingerprints. As desired, both predictions are highly correlated (PCC ≈ 1) and the mean absolute difference between them is very small (0.8). Figure 5B shows the same analysis for a model where we removed the duplicated entries from the training data. Even though the correlation is still high (PCC = 0.82), it decreased strongly, while prediction differences increased notably to 9.12 on average. The mean PCCs per drug (for monotherapies) and per drug combination are 0.98 and 0.97 for the duplicated training data and decrease to 0.78 and 0.86 for the non-duplicated training data, respectively. This is also represented in the test error where the model with duplicated training entries achieved an MAE of 12.39 compared to 14.6 for non-duplicated entries. Similar trends can also be observed for the PhysChem setting (see Supplementary Figure 5).

**Figure 5:**
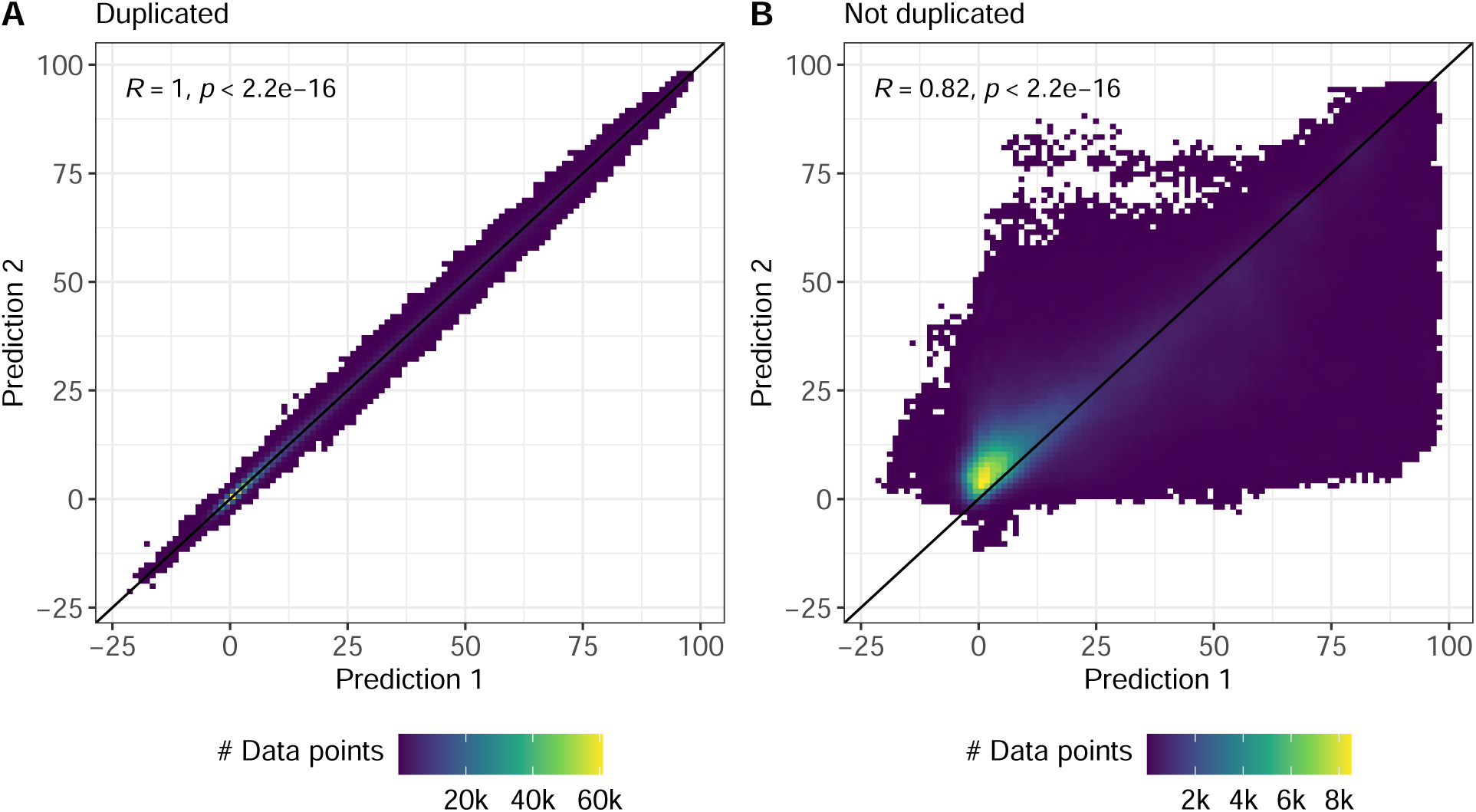
Correlation of duplicated entries from the test data. This figure shows the correlation between the model predictions for duplicated entries. Duplicated entries refer to the same drug-drug-cell combination and the same treatment concentrations but can be represented by two different model inputs through swapping the features of the respective drugs (cf. Methods and Figure 1) Sub-figure A shows the test predictions when including duplicated entries into the training data, while Sub-figure B shows the predictions when training only on non-duplicated entries. In both figures, the black diagonal line represents the identity and R denotes the Pearson correlation between the predictions.

### Reconstruction of Drug Sensitivity Measures

A benefit of predicting concentration-specific inhibition values is that based on the model’s predictions, dose-response curves and matrices can be reconstructed. These can in turn be used to compute various measures of drug sensitivity or synergy. Since the focus of this paper is on sensitivity prediction and Vlot et al. discourage the computation of arbitrary synergy scores on large-scale data,^13^ we reconstructed two measures of drug sensitivity, namely our recently published measure called *CMax viability* for monotherapies, and a modification of this measure for drug combinations, which we call the *combination CMax viability* (cf. Methods). Unlike conventional sensitivity measures like the IC50 or AUC, the (combination) CMax viability is comparable across drugs^15^ and drug combinations. Consequently, it can be used to prioritize drugs/combinations for a given cell line (i.e., rank them by their effectiveness), which will be investigated in the next section.

For the computation of monotherapy CMax viabilities, we first used the actual/predicted monotherapy entries of the test data to generate actual/predicted dose-response-curves (cf. Methods). An example is shown in Figure 2, where we also highlight how the CMax viability is derived from the curves. In total, we were able to compute both the actual and predicted CMax viabilities for 7,352 out of 32,564 cell line-drug combinations. The decreased number of combinations stems from the fact that CMax concentrations were only available for 77 of the investigated drugs. Figure 6 depicts the prediction errors for the reconstructed monotherapy CMax viability values. The mean MAE averaged over all drugs is 0.12 and the mean MSE is 0.04, which is comparable to the error we previously achieved when predicting CMax viabilities directly using either the SAURON-RF algorithm by Lenhof et al. ^29^ (MSE = 0.03) or a slightly adjusted version of DeepDR by Chiu et al. ^54^ (MSE = 0.09). ^15^ A baseline error can be obtained from a model that for every treatment concentration predicts the mean inhibition for each drug obtained from the training data. For such a model, the CMax viability (i.e., the viability at the CMax concentration) would also be predicted as this mean. This would result in a baseline MAE of 0.2, which our model improves by 40%. The overall PCC is 0.58 for the CMax viabilities and 0.41 for the baseline. However, the drug-specific PCC is only 0.1 (cf. Figure 6B). While a drug-specific baseline PCC cannot be computed for constant predictions, adding random noise with mean 0 to these constant predictions results in a baseline PCC of 0. Thus, our predictions improve this baseline but only slightly. When using our models to reconstruct IC50 values, we observe a similar phenomenon (overall PCC = 0.71, mean PCC per drug = 0.01, cf. Supplementary Figure 6). To investigate the reasons for these low drug-specific correlations, we developed and evaluated different hypotheses, which can be found in the Supplement. Based on our evaluation of these hypotheses, we conclude that even though prediction errors are relatively small and comparable to our previous work, the derived measures cannot be used to compare the effect of a drug monotherapy on different cell lines. For the combination CMax viability (26,946 drug-drug-cell line combinations), we obtained similar results, which are depicted in Figure 6C and D.

**Figure 6:**
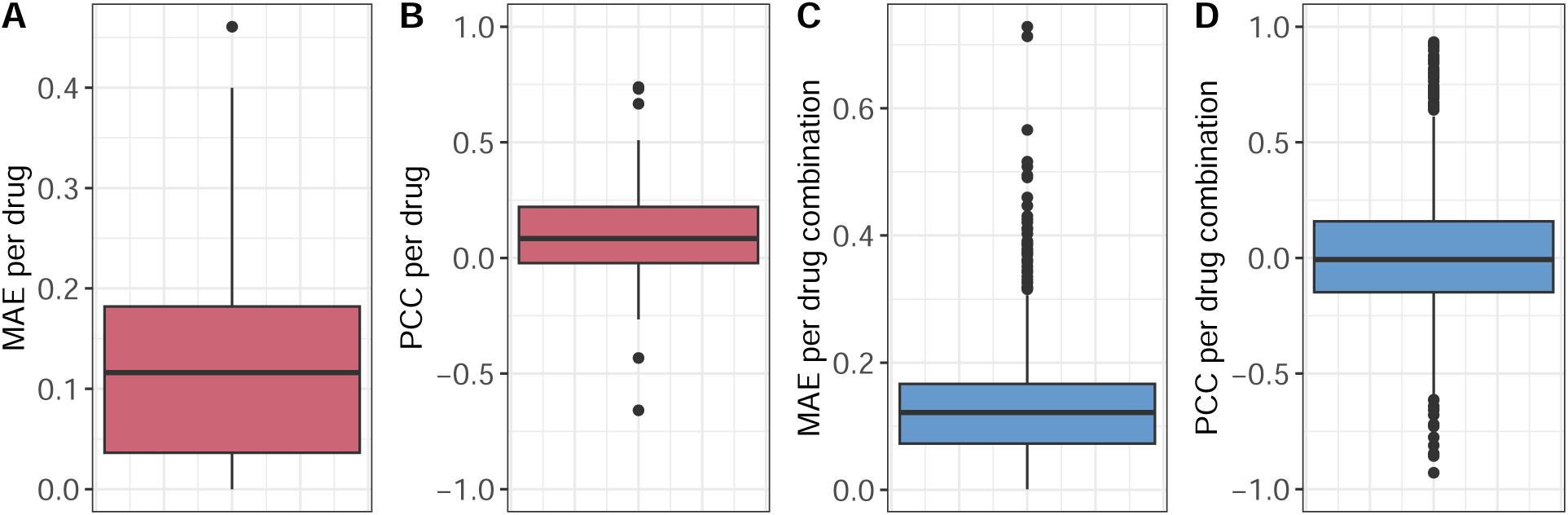
Reconstruction of (combination) CMax viabilities from predicted dose-response curves/matrices. Sub-figures A and B (red) show the distribution of MAE and PCC per drug for the reconstruction of CMax viabilities using dose-response curves fit on the test set monotherapy data. Sub-figures C and D (blue) show the distribution of MAE and PCC per dug combination for the reconstruction of combination CMax viabilities using dose-response matrices derived from the test set drug combination data.

Nevertheless, we would like to highlight that such an evaluation of drug-specific correlations as conducted here is frequently not performed for drug sensitivity and synergy prediction (cf. Supplementary Table 3, where we compare the investigated settings and analyses for 39 state-of-the-art methods). Thus, similar problems may often go undetected.

Due to the novelty of our prediction approach, there is no method we could directly compare our findings to. Nevertheless, our analyses presented earlier show that our models are competitive in performance to the approaches by Zheng et al. ^5^ and Julkunen et al. ^18^ for making predictions using known cell lines and drug combination data. Note that both approaches do not provide drug-/combination-specific correlations.

For cell-blind evaluations on monotherapy data, we found three related approaches that provide drug-specific correlations: Our recently published method SAURON-RF achieves a mean PCC of 0.56 when directly predicting CMax viabilities using drug-specific models. ^15^ In the same manuscript we also show that an adjusted version of the multi-drug model DeepDR by Chiu et al. ^54^ achieves a PCC of 0 for the same task. In comparison, Chawla et al. employ multi-drug models for the prediction of IC50 values and achieve mean PCCs between ca. 0.18 and 0.5 for different ML algorithms. Lastly, Rahman and Pal achieve mean PCCs between 0.29 and 0.44 when reconstructing AUC values from predicted dose-response curves. While not directly comparable to our approach, these works underline that at least weak to moderate drug-specific correlations can be achieved (1) for predicting CMax viabilities (2) when using multi-drug models (3) when deriving sensitivity measures from predicted curves. Yet, it remains to be investigated further if and how comparable results can be achieved when combining all three factors and also considering combination therapies, thereby enabling predictions for arbitrary drugs/combinations and measures, which we aim to achieve here.

### Treatment Prioritization

In our final analysis, we investigate how accurately drugs and drug combinations can be prioritized for a given cell line based on the model predictions: For each cell line in the test set, we used the computed CMax viabilities for the monotherapy and combination data to achieve a ranking of drugs and drug combinations from most to least effective. Drug prioritization is supposed to mimic a personalized treatment scenario with the goal to achieve a list of most effective treatment suggestions for a given patient. The results are shown in Figure 7, where the first row shows the results for monotherapies only, while the second row shows the results when combining mono- and combination therapies into one list. The results for combination therapies only are shown in Supplementary Figure 8.

**Figure 7:**
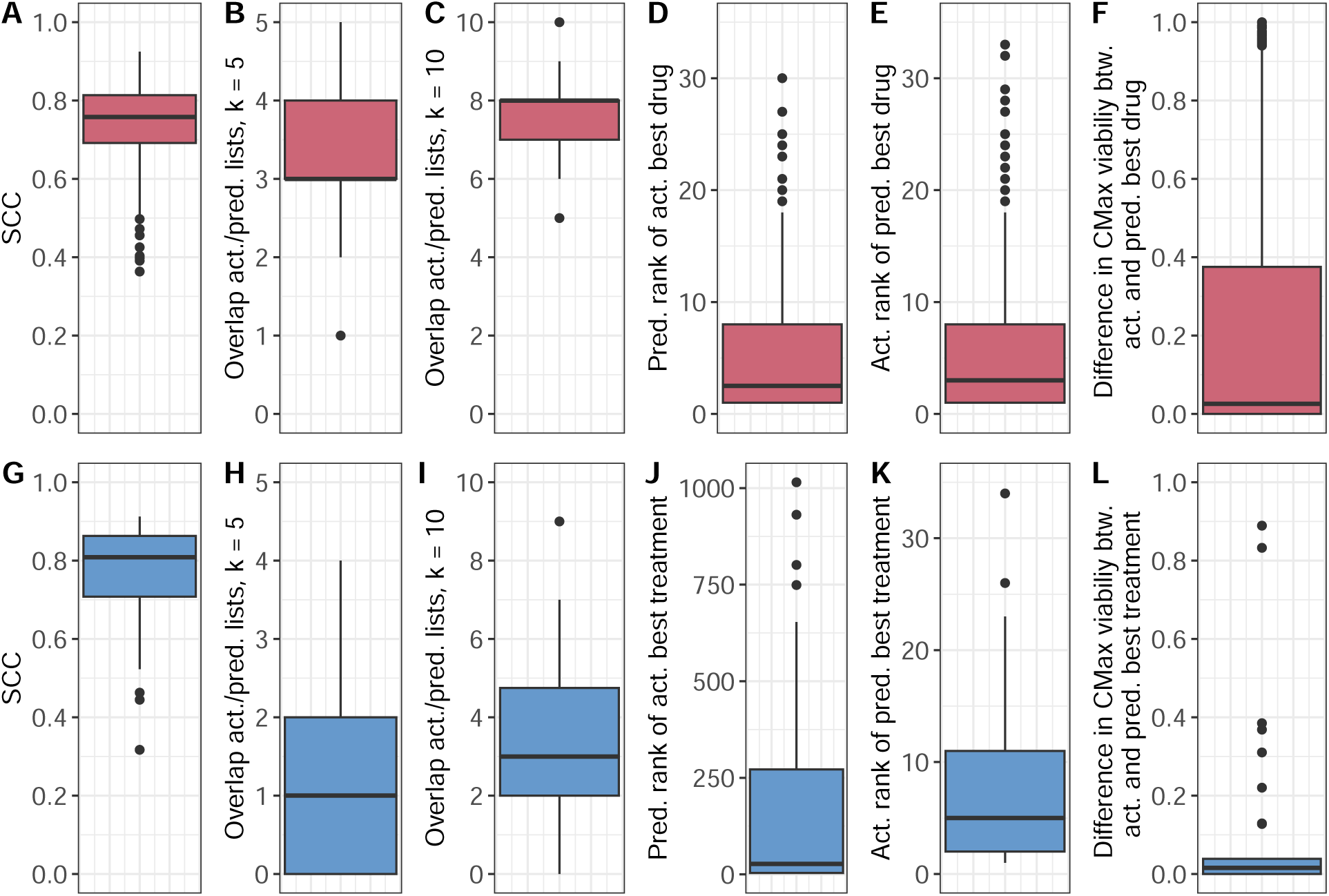
Treatment prioritization. This figure depicts the test set prioritization results for mono- and combination therapies. Sub-figures A to F (red) focus on the prioritization of monotherapies including: (A) the SCC between the actual and predicted rankings for each cell line, (B)/(C) the intersection size between the 5/10 actual and predicted most effective treatments, (D) the predicted rank of the actual most effective treatment, (E) the actual rank of the treatment predicted to be most effective, and (F) the difference between the actual CMax viabilities for the actual and predicted most effective treatment. Sub-figures G to L (blue) show the analogous prioritization results when combining mono- and combination treatments into one list.

For monotherapies, the Spearman correlation coefficient (SCC) between the actual and predicted rankings was 0.74 (baseline (as defined in the previous section): 0.54). Our predictions clearly outperform the baseline. Still, the baseline correlation is relatively high, indicating that the differences in effectiveness between drugs are easier to predict than the differences between cell lines receiving the same treatment.

While an accurate ranking for the entire list is desirable, one would typically place more emphasis on the correct identification of the most effective treatments. Thus, we computed the mean overlap between the first *k* elements of the actual and predicted rankings. For monotherapies, the average length of the predicted drug lists is 31.15. The average overlap between the top *k* = 5 and *k* = 10 actual and predicted most effective drugs is 3.16 (baseline: 2.14) and 7.68 (baseline: 6.55), respectively (results for further *k* are shown in Supplementary Figure 9). Furthermore, the median rank of the actually most effective drug in the predicted ranking is 2.5 (baseline: 8), and the median rank of the drug predicted to be most effective in the actual list is 3 (baseline 6). The median difference between the true CMax viabilities of the actual most effective and predicted most effective drugs is only 0.02 (baseline 0.31).

The second row of Figure 7 shows the analogous prioritization results when combining mono- and combination treatments into one list. The SCC of 0.76 (baseline: 0.62) is comparable to the results for monotherapies. Since the average list length is much greater when including drug combinations (838.62), the overlaps at *k* = 5 (1.26, baseline: 0.68) and *k* = 10 (3.38, baseline: 2.09) are lower (cf. also Supplementary Figures 9 and 10). Furthermore, the median rank of the actually best treatment in the predicted list (27, baseline: 170.5) and of the predicted best treatment in the actual list (9.5, baseline: 12) decrease. Still, results clearly improve over the baseline. Furthermore, the median difference in viability between the actually most effective treatment and the treatment predicted to be most effective remains small (0.02, baseline 0.03).

## Discussion

Administering not only single but multiple drugs in combination is common in cancer treatment. However, while drug response datasets for monotherapy data have been available for more than a decade, large-scale data sets for combination therapy have only become publicly available more recently, e.g., the DrugComb database. ^4,5^ While the DrugComb data have extensively been studied for the prediction of drug synergy, they are still underused for the prediction of drug sensitivity, especially with the focus on making personalized treatment recommendations. For this application case, we found the scores that are widely used for synergy prediction less suited due to various reasons discussed in this manuscript.

To exploit the available drug combination data for predicting drug responses without relying on synergy scores, we developed and evaluated several ML algorithms and architectures that directly predict concentration-specific drug responses in the form of relative inhibitions. We are convinced that this approach has various benefits for personalized treatment recommendation: First, our approach allows the reconstruction of dose-response curves and matrices from the model predictions. From these curves/matrices, various sensitivity or synergy measures can be reconstructed. The inspection of individual curves/matrices can aid in validating the underlying assumptions for certain measures. Next, our approach can predict both mono- and combination therapies. Additionally, our approach allows for making predictions for unseen cell lines, thereby mimicking the scenario assessing drug responses for a new patient. Together with our novel sensitivity measure, the *(combination) CMax viability*, this framework finally enables the prioritization of both mono- and combination therapy options for unseen cell lines (patients).

Our evaluations on the DrugComb database show that our models substantially improve baseline models and show very little variation when predicting the same treatment using different input representations. Notably, we evaluated our models on unseen cell lines, which is often neglected in drug sensitivity prediction. ^19^ Moreover, our models are also competitive with state-of-the-art approaches when making predictions for known cell lines. Furthermore, we achieved strong correlations for treatment prioritization. However, our analyses also reveal weaknesses of directly predicting relative inhibitions: While prediction errors for the reconstruction of drug response measures are competitive with other approaches, the drug-specific correlations between these measures only slightly improve over a baseline model. Additionally, we observed increased prediction errors for data samples with high inhibitions, corresponding to cases of treatment sensitivity. This issue is relatively well-known for classification but has rarely been discussed or addressed for regression. ^29,34^

Three main factors can be adjusted to potentially address such challenges, namely the choice of ML algorithm, the choice and representation of input features, and the used data:

**ML algorithm:** We investigated neural networks (highly popular for sensitivity and synergy prediction), random forests, and elastic nets. In our recently published benchmarking, we found both elastic nets and random forests to outperform neural networks when predicting drug sensitivity.^24^ For the prediction of inhibitions, as investigated here, random forests are superior to the other algorithms. In general, a plethora of further (potentially more sophisticated) approaches can be used to model the prediction of inhibitions. However, as discussed in our benchmarking^24^ and also by Li et al., ^55^ more complex approaches are not necessarily superior to simpler ML algorithms, and careful evaluation is required to ensure a fair performance comparison.

**Input features and representation:** For the characterization of cell lines in the model input, several sources found gene expression to be the most informative omics-type for predicting drug responses. ^54,56,57^ However, the inclusion of further omics or a priori knowledge, e.g., known sensitivity biomarkers or protein interactions, might improve predictions.

Similarly, further drug properties, e.g., Morgan fingerprints^58^ could be investigated, or graph neural networks could be employed to represent drugs as molecular graphs. However, the superiority of molecular graphs over conventional drug fingerprints for sensitivity/synergy prediction and drug discovery has been questioned. ^27,59^

**Dataset:** With 947 cell lines, 265 drugs, and 9,535 drug combinations, the dataset invested here is notably larger compared to other approaches working on drug combination data.^5,12,14,18,49,60,61^ Unfortunately, given the size of the investigated dataset, hardware restrictions become a limiting factor for ML. Despite training models on a compute cluster with machines of 500 gigabytes working memory, we had to reduce our data regarding the number of considered drugs, features, and methods (cf. Methods).

Generally, a large amount of training data benefits model training and robustness. Yet, if the dataset is heterogeneous, e.g., due to different data sources, as is the case for DrugComb, this may decrease performance compared to models built and evaluated on a more homogeneous dataset. Even though Zagidullin et al. found the reproducibility between replicates from different datasets satisfactory in the first release of DrugComb,^4^ disagreement between drug response data from different sources is a well-known problem. ^57,62,63^ Especially for clinical applications, combining data from different sources (e.g., different hospitals) is essential, and models should be able to cope with this degree of heterogeneity. To this end, meta- or transfer-learning methods could be leveraged. ^64^

Investigating different ML algorithms, input representations, and datasets can potentially improve the predictive performance. However, especially in a sensitive field such as personalized medicine, performance alone should not be regarded the sole building block of model trustworthiness. ^65^ E.g., to assess the reliability of individual predictions, uncertainty estimation frameworks like conformal prediction could be applied. ^15,66–68^ Additionally, incorporating interpretability mechanisms^12,65,69^ into the model design and evaluation can aid in identifying drug or cell line properties that impact the predicted response. This could not only make predictions more comprehensible but also be useful to infer novel mechanisms of drug sensitivity or synergy.

## Data and Software Availability

The drug response data used for our analyses can be downloaded from the DrugComb website (https://drugcomb.org/download) and the DrugComb API (https://api.drugcomb.org/, cf. Methods). The gene expression data of canncer cell lines can be downloaded from the GDSC website (https://www.cancerrxgene.org/downloads/bulk_download). CMax concentrations for 77 of the investigated drugs can be derived from Liston and Davis. ^20^ Our code is available at GitHub (https://github.com/unisb-bioinf/Drug_Combination_Sensitivity_Prediction), where we also provide the SMILES, MACCS fingerprints and physico-chemical properties derived from RDKit,^21^ as well as the one-hot encoded target molecules of the investigated compounds.

