## Supplement for "How to Predict Effective Drug Combinations - Moving beyond Synergy Scores"

#### 1 Definition of Relative Inhibition

Typically, cell viability assays measure the presence of live cells through fluorescence/luminescence intensities [1, 2, 3]. These intensities are further processed to obtain relative inhibitions. To this end, a background correction is performed by subtracting the intensity of the positive control wells (i.e., wells with only medium but no cells) from the intensity of the negative control wells (i.e., wells with untreated cells and medium) and all treatment wells (i.e., wells with treated cells and medium) [4].
Next, relative viabilities are computed by dividing the background-corrected intensities of all treatment wells with the negative control [4]. To obtain relative inhibitions, we subtract the relative viability from 1. In the DrugComb database, the resulting values are additionally multiplied by 100.
Typically, relative inhibitions are in range $(-\infty, 100]$, where values $< 0$ indicate that the treatment increases cell growth and values $> 0$ indicate a reduction in growth. While values $> 100$ should generally not occur, it can happen that the background correction becomes negative if the intensity of the treatment well is smaller than that of the control in cases where the treatment killed all cells. Note that Zheng et al., i.e., the authors of DrugComb, recommend considering only relative inhibitions in $[-200, 200]$ as data points outside of this range are deemed to be of poor quality. In contrast, the Genomics of Drug Sensitivity in Cancer database limits the considered intervals more strictly [5].

#### 2 Synergy Scores

In this section, we give a brief overview on four commonly used synergy scores, namely the HSA [6], Bliss [7], Loewe [8], and ZIP [9] synergy score. Our descriptions are limited to experiments using two-drug combinations. Extensions for an arbitrary number of drugs are provided at [10].

Consider an experiment where a cell line $c$ is treated with different concentrations of two drugs, $d_1$ and $d_2$. For $d_1$, $n$ different concentrations were tested, for $d_2$, $m$ different concentrations were tested. The results of the combination treatments are reported in an $n \times m$ dose-response matrix $Y$, where each entry $y_{a,b}$ denotes the percentage of relative inhibition after administering dose $a$ of $d_1$ in combination with dose $b$ of $d_2$. Additionally, monotherapy responses for the same $n$ ($m$) concentrations are required. We denote the percentage of relative inhibition obtained from the monotherapy of $d_1$ with concentration $a$ as $y_a$. Analogously, the percentage of relative inhibition obtained from the monotherapy of $d_2$ with concentration $b$ is given by $y_b$.
To measure the synergy between $d_1$ and $d_2$ on cell line $c$, the observed inhibitions $y_{a,b}$ are compared to the estimated inhibitions $\hat{y}_{a,b}$ that are calculated using a reference model which assumes no synergistic or antagonistic interaction between the two drugs. For the DrugComb database, synergy scores were computed using the widely applied synergyfinder R package [11], which calculates the synergy score ($SS$) for a cell-drug-drug combination as the average over all concentration-specific comparisons between the observed and the estimated drug responses [10]:

$$SS = \frac{1}{n \cdot m} \sum_{a \in \mathcal{A}} \sum_{b \in \mathcal{B}} (y_{a,b} - \hat{y}_{a,b}) \qquad (1)$$

Here, $\mathcal{A}$ and $\mathcal{B}$ denote the sets containing all tested doses for drug $d_1$ and $d_2$, respectively. For $SS > 0$, the observed inhibitions are on average greater than the expected inhibitions, indicating synergy between $d_1$ and $d_2$. In contrast, for $SS < 0$, the observed inhibitions are smaller than expected, indicating an antagonistic interaction between the drugs.
Note that the synergyfinder documentation states that the percentage relative inhibitions utilized to derive $SS$ are typically expected to be in range $[0, 1]$, while values outside of this range are still allowed [12]. While there is a (non-default) option to baseline-correct negative inhibitions [12], we did not find documentation on how synergyfinder handles negative values or values greater than 1 when present.

In the following, we present four of the most common reference models for estimating $\hat{y}_{a,b}$.

**HSA [6]:** The highest single agent (HSA) model expects the effect of a non-interacting drug combination to be equal to the greater of both monotherapy effects at the same concentrations [6]:

$$\hat{y}_{a,b}^{HSA} = max(y_a, y_b) \tag{2}$$

$$\tag{3}$$

**Bliss [7]:** The Bliss model assumes the combination effect of two drugs to be statistically independent [7]:

$$\hat{y}_{a,b}^{Bliss} = y_a + y_b - y_a \cdot y_b \tag{4}$$

**Loewe [8]:** The Loewe model assumes that there exists a concentration $A$ of $d_1$ and a concentration $B$ of $d_2$, for which the monotherapy can achieve the same effect as the combination of both drugs:

$$y_{a,b} = y_A = y_B \tag{5}$$

Additionally, it is assumed that for each concentration of drug $d_1$, there exists a concentration of drug $d_2$ with the same effect and vice versa. Consequently, we can define

$$A = a + a_b \tag{6}$$

$$B = b + b_a \quad , \tag{7}$$

where, $a_b$ denotes the concentration of $d_1$ for which $y_{a_b} = y_b$. Analogously, $b_a$ denotes the concentration of $d_2$ for which $y_{b_a} = y_a$.
The model furthermore assumes that the potency ratio between the two drugs is constant over the entire dose-response curve. This means the ratio between two concentrations from $d_1$ and $d_2$ achieving the same effect is a constant $R$. In particular, it holds that

$$R = \frac{A}{B} = \frac{a_b}{b} = \frac{a}{b_a} \quad . \tag{8}$$

It follows that the expected combination response exhibits so-called *Loewe additivity* [9]:

$$a + a_b = A \tag{9}$$

$$\Leftrightarrow a + b \cdot R = A \tag{10}$$

$$\Leftrightarrow a + b \cdot \frac{A}{B} = A \tag{11}$$

$$\Leftrightarrow \frac{a}{A} + \frac{b}{B} = 1 \tag{12}$$

To identify $A$ and $B$, and, consequently, to compute $\hat{y}_{a,b}^{Loewe}$, dose-response curves for both drugs need to be fit, which are generally modeled as four-parametric logistic functions [9]. We omit the details here but refer interested readers to Yadav et al. for a detailed description [9], which concludes that $\hat{y}_{a,b}^{Loewe}$ can finally be computed by solving:

$$\frac{a}{m_1 \cdot \left(\frac{\hat{y}_{a,b}^{Loewe} - E_{min}^1}{E_{max}^1 - \hat{y}_{a,b}^{Loewe}}\right)^{\frac{1}{\lambda_1}}} + \frac{b}{m_2 \cdot \left(\frac{\hat{y}_{a,b}^{Loewe} - E_{min}^2}{E_{max}^1 - \hat{y}_{a,b}^{Loewe}}\right)^{\frac{1}{\lambda_2}}} = 1 \tag{13}$$

Here, $E^1_{min}$ and $E^1_{max}$ are derived from the fitted dose-response curve and denote the minimum and maximum effect that drug $d_1$ can achieve. Additionally, $\lambda_1$ denotes the slope and $m_1$ the midpoint of the dose-response curve for $d_1$. $E^2_{min}$, $E^2_{max}$, $\lambda_2$ and $m_2$ are defined analogously for drug $d_2$.

**ZIP [9]:** The *zero interaction potency* (ZIP) model combines ideas of both the Bliss and Loewe models. It models the notion of non-interaction between drugs by assuming that the dose-response curve of one drug is unaffected by the addition of the second drug. Consequently, the combination effect can be described by shifting the dose-response curve of either drug by the effect of the other. Shifting the curve of drug $d_1$ by the effect of drug $d_2$ at concentration $b$ can be described as follows (assuming that $E^1_{min} = 0$ and $E^1_{max} = 1$) [9]:

$$\hat{y}_{1\leftarrow 2} = \frac{y_b + \frac{x}{m_1}{}^{\lambda}_{1}}{1 + (\frac{x}{m_1})^{\lambda}_{1}} \tag{14}$$

Here, $x$ denotes any dose of $d_1$. Thus, the conventional dose-response curve for $d_1$ is modified by simply adding $y_b$ in the numerator, thereby increasing the baseline effect. Analogously, $\hat{y}_{2\leftarrow 1}$ can be defined. Consequently, just like the Loewe score, the ZIP score relies on accurate curve fittings for both monotherapies. Yadav et al. show that both $\hat{y}_{1\leftarrow 2}$ and $\hat{y}_{2\leftarrow 1}$ are equivalent to estimating the combined drug response as follows [9]:

$$\hat{y}^{ZIP}_{a,b} = \frac{\left(\frac{a}{m_1}\right)^{\lambda_1}}{1 + \left(\frac{a}{m_1}\right)^{\lambda_1}} + \frac{\left(\frac{b}{m_2}\right)^{\lambda_2}}{1 + \left(\frac{b}{m_2}\right)^{\lambda_2}} - \frac{\left(\frac{a}{m_1}\right)^{\lambda_1}}{1 + \left(\frac{a}{m_1}\right)^{\lambda_1}} \cdot \frac{\left(\frac{b}{m_2}\right)^{\lambda_2}}{1 + \left(\frac{b}{m_2}\right)^{\lambda_2}} \tag{15}$$

Similar to the Bliss score provided in Equation 4, Equation 15 also follows the form $X + Y - X \cdot Y$. Consequently, a non-interaction in the ZIP model ($\hat{y}^{ZIP}_{a,b} = 0$) also corresponds to probabilistic independence [9].

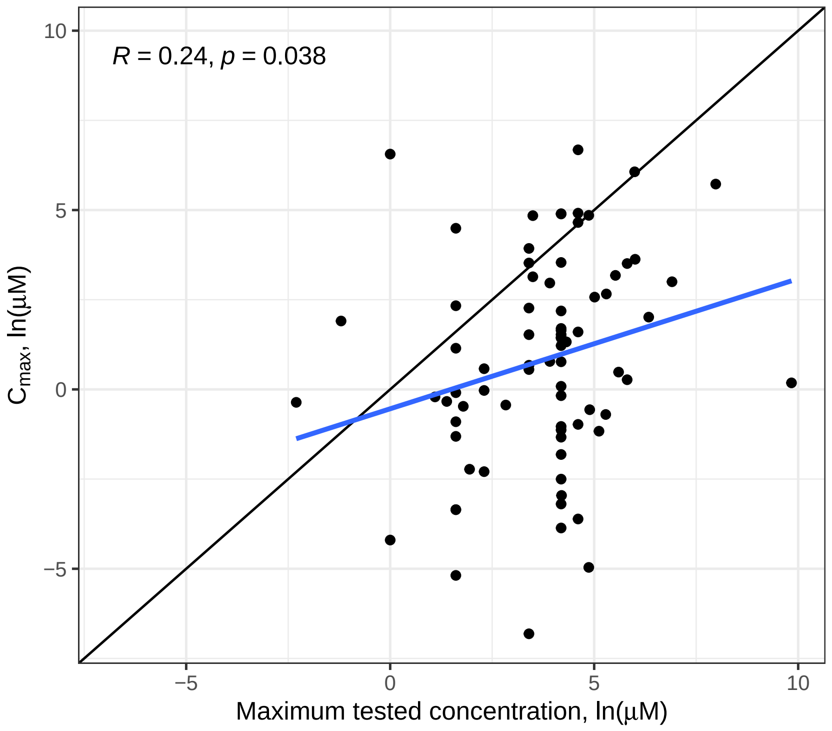


Figure 1: Comparison of CMax values and maximum tested drug concentrations. This figure depicts the CMax concentrations for 77 drugs from DrugComb in comparison to the maximum screened concentrations in our investigated dataset. The CMax concentrations were obtained from [13] and denote the peak plasma concentration of a drug after administering the highest clinically recommended dose. Additionally, the Pearson correlation coefficient (R) and a regression line (blue) are depicted.

Table 1: Investigated Compounds. Table continues on next pages.

| | Compound Name / CAS Number | | Compound Name / CAS Number |
|---|---|---|---|
| 1 | Abiraterone | 51 | CHIR-99021 |
| 2 | actinomycin D | 52 | chlorambucil |
| 3 | ADM hydrochloride | 53 | cis-Platin |
| 4 | Afatinib | 54 | Co-V |
| 5 | Akt inhibitor VIII | 55 | Crizotinib |
| 6 | allopurinol | 56 | cyclophosphamide |
| 7 | alpelisib | 57 | Cylocide |
| 8 | altretamine | 58 | CYTARABINE HYDROCHLORIDE |
| 9 | amifostine | 59 | dacarbazine |
| 10 | anastrozole | 60 | Daporinad |
| 11 | Antibiotic AD 32 | 61 | Darinaparsin |
| 12 | Antibiotic AY 22989 | 62 | Dasatinib |
| 13 | Avagacestat | 63 | Decitabine |
| 14 | Axitinib | 64 | Deforolimus |
| 15 | Azacytidine, 5- | 65 | dexamethasone |
| 16 | AZD1208 | 66 | Dexrazoxane |
| 17 | AZD1480 | 67 | Dinaciclib |
| 18 | AZD2014 | 68 | docetaxel |
| 19 | AZD4320 | 69 | Doramapimod |
| 20 | AZD4547 | 70 | dorsomorphin |
| 21 | AZD5363 | 71 | Dovitinib |
| 22 | AZD5582 | 72 | doxorubicin |
| 23 | AZD6482 | 73 | Elesclomol |
| 24 | AZD6738 | 74 | Eloxatin (TN) |
| 25 | AZD7762 | 75 | Eloxatin (TN) (Sanofi Synthelab) |
| 26 | AZD8055 | 76 | EMBELIN |
| 27 | AZD8186 | 77 | Emcyt (Pharmacia) |
| 28 | Belinostat | 78 | Entinostat |
| 29 | Bendamustine hydrochloride | 79 | Enzastaurin |
| 30 | Bexarotene | 80 | Erlotinib |
| 31 | BI-78D3 | 81 | Erlotinib hydrochloride |
| 32 | BI-D1870 | 82 | etoposide |
| 33 | BI 2536 | 83 | EXEMESTANE |
| 34 | bicalutamide | 84 | Fedratinib |
| 35 | Bleo | 85 | FH535 |
| 36 | bleomycin | 86 | Fingolimod |
| 37 | BMS-536924 | 87 | Fludarabine Base |
| 38 | BMS-754807 | 88 | Foretinib |
| 39 | Bortezomib | 89 | Fulvestrant |
| 40 | Bosutinib | 90 | GDC-0879 |
| 41 | busulfan | 91 | Gefitinib |
| 42 | CABAZITAXEL | 92 | geldanamycin |
| 43 | Cabozantinib | 93 | gemcitabine |
| 44 | Carboplatinum | 94 | GSK 650394 |
| 45 | carmustine | 95 | GSK429286A |
| 46 | Cediranib | 96 | GW 441756 |
| 47 | celecoxib | 97 | GW0742 |
| 48 | CHEMBL17639 | 98 | GW2580 |
| 49 | CHEMBL277800 | 99 | GW843682X |
| 50 | CHEMBL3103192 | 100 | hydroxyurea |

Continuation of Table 1

| | Compound Name / CAS Number | | Compound Name / CAS Number |
|---|---|---|---|
| 101 | Idelalisib | 151 | OSI-027 |
| 102 | ifosfamide | 152 | OSI-930 |
| 103 | Imatinib | 153 | OSU-03012 |
| 104 | IMD-0354 | 154 | paclitaxel |
| 105 | IMIQUIMOD | 155 | Palbociclib |
| 106 | IPA-3 | 156 | Panobinostat |
| 107 | Ixabepilone | 157 | parthenolide |
| 108 | JZL184 | 158 | Pazopanib |
| 109 | Ku-0063794 | 159 | Pazopanib hydrochloride |
| 110 | KU-55933 | 160 | Pemetrexed |
| 111 | KU-60019 | 161 | Perifosine |
| 112 | l-685,458 | 162 | PF-04217903 |
| 113 | L-778123 free base | 163 | PF-562271 |
| 114 | Lapatinib | 164 | PHA-793887 |
| 115 | Lenalidomide | 165 | PI-103 |
| 116 | Lestaurtinib | 166 | PIK-93 |
| 117 | letrozole | 167 | Pioglitazone |
| 118 | lfm-a13 | 168 | Piperlongumine |
| 119 | Linifanib | 169 | pipobroman |
| 120 | Linsitinib | 170 | PLX-4720 |
| 121 | lomustine | 171 | Pralatrexate |
| 122 | Masitinib | 172 | Procarbazine hydrochloride |
| 123 | MEGESTROL ACETATE | 173 | QS11 |
| 124 | Melphalan hydrochloride | 174 | Quinacrine hydrochloride |
| 125 | metformin | 175 | Quizartinib |
| 126 | methotrexate | 176 | RAF265 |
| 127 | methoxsalen | 177 | raloxifene |
| 128 | Midostaurin | 178 | Retinoic acid |
| 129 | MITHRAMYCIN | 179 | Romidepsin |
| 130 | mitomycin C | 180 | Ruxolitinib |
| 131 | mitotane | 181 | Sapitinib |
| 132 | mitoxantrone | 182 | Saracatinib |
| 133 | MK-1775 | 183 | Selumetinib |
| 134 | MK-2206 | 184 | Serdemetan |
| 135 | MK-4541 | 185 | Silmitasertib |
| 136 | MK-5108 | 186 | SNS-032 |
| 137 | MLN4924 | 187 | SNX-2112 |
| 138 | MRK003 | 188 | Sorafenib |
| 139 | Navelbine ditartrate (TN) | 189 | Sunitinib |
| 140 | Navitoclax | 190 | T0901317 |
| 141 | Nilotinib | 191 | Tamoxan |
| 142 | Niraparib | 192 | Tamoxifen citrate |
| 143 | NSC-127716 | 193 | Tanespimycin |
| 144 | NSC256439 | 194 | temozolomide |
| 145 | NSC609699 | 195 | Temsirolimus |
| 146 | NSC733504 | 196 | teniposide |
| 147 | NSC756645 | 197 | TGX-221 |
| 148 | Nutlin-3 | 198 | thalidomide |
| 149 | Olaparib | 199 | thapsigargin |
| 150 | Onalespib | 200 | thiotepa |

Continuation of Table 1

| | Compound Name / CAS Number |
|---|---|
| 201 | Tipifarnib |
| 202 | Tivozanib |
| 203 | topotecan |
| 204 | TOPOTECAN HYDROCHLORIDE |
| 205 | Tozasertib |
| 206 | TPCA-1 |
| 207 | Trametinib |
| 208 | Triethylenemelamine |
| 209 | Trisenox |
| 210 | Tubastatin A |
| 211 | TW-37 |
| 212 | UNC0638 |
| 213 | Uramustine |
| 214 | US9505780, JQ-1 |
| 215 | Vandetanib |
| 216 | Veliparib |
| 217 | Vemurafenib |
| 218 | Vepesid J |
| 219 | vinblastine |
| 220 | Vinblastine sulfate |
| 221 | vincristine |
| 222 | Vincristine sulfate |
| 223 | vinorelbine |
| 224 | Vismodegib |
| 225 | Vorinostat |
| 226 | VX-702 |
| 227 | XL147 |
| 228 | XL765 |
| 229 | YK 4-279 |
| 230 | Zanosar |
| 231 | Zoledronic acid |
| 232 | ZSTK474 |
| 233 | (-)-Rapamycin |
| 234 | 001, RAD |
| 235 | 1032350-13-2 |
| 236 | 122111-05-1 |
| 237 | 1260907-17-2 |
| 238 | 158798-73-3 |
| 239 | 218137-86-1 |
| 240 | 219580-11-7 |
| 241 | 23541-50-6 |
| 242 | 284028-89-3 |
| 243 | 303727-31-3 |
| 244 | 315183-21-2 |
| 245 | 391210-10-9 |
| 246 | 49843-98-3 |
| 247 | 5-Aminolevulinic acid hydrochloride |
| 248 | 5-azacytidine |
| 249 | 5-Fluoro-2'-deoxyuridine |
| 250 | 5-Fluorouracil |

| | Compound Name / CAS Number |
|---|---|
| 251 | 547757-23-3 |
| 252 | 55-86-7 |
| 253 | 6-Mercaptopurine |
| 254 | 6-Thioguanine |
| 255 | 7-Ethyl-10-hydroxycamptothecin |
| 256 | 717906-29-1 |
| 257 | 761439-42-3 |
| 258 | 7803-88-5 |
| 259 | 781661-94-7 |
| 260 | 803712-79-0 |
| 261 | 841290-80-0 |
| 262 | 844499-71-4 |
| 263 | 891494-63-6 |
| 264 | 915019-65-7 |
| 265 | 957054-30-7 |

Table 2: This table denotes the tuned hyperparameters for each ML algorithm and setting. For hyperparameters not stated explicitly, the values denoted in Table 1 of the main manuscript were employed. Otherwise, we used the default parameters as provided the respective Python packages.

| Model | Setting | Parameters |
|---|---|---|
| Random Forest | OneHot | max_features=250; max_depth=1000000; min_samples_leaf=2 |
| Random Forest | OneHotTar | max_features=250; max_depth=1000000; min_samples_leaf=2 |
| Random Forest | MACCS | max_features=250; max_depth=100; min_samples_leaf=2 |
| Random Forest | PhysChem | max_features=100; max_depth=100; min_samples_leaf=2 |
| Neural Network | OneHot | activation=elu; learning_rate=0.0001; num_hidden_layers=5; dropout=0.3; |
| Neural Network | OneHotTar | activation=elu; learning_rate=0.0001; num_hidden_layers=4; dropout=0.3; |
| Neural Network | MACCS | activation=elu; learning_rate=0.0001; num_hidden_layers=5; dropout=0.3; |
| Neural Network | PhysChem | activation=tanh; learning_rate=0.0001; num_hidden_layers=4; dropout=0.1; |
| Elastic Net | OneHot | alpha=0.01; l1_ratio=1 |
| Elastic Net | OneHotTar | alpha=0.01; l1_ratio=1 |
| Elastic Net | MACCS | alpha=0.01; l1_ratio=1 |
| Elastic Net | PhysChem | alpha=0.01; l1_ratio=1 |

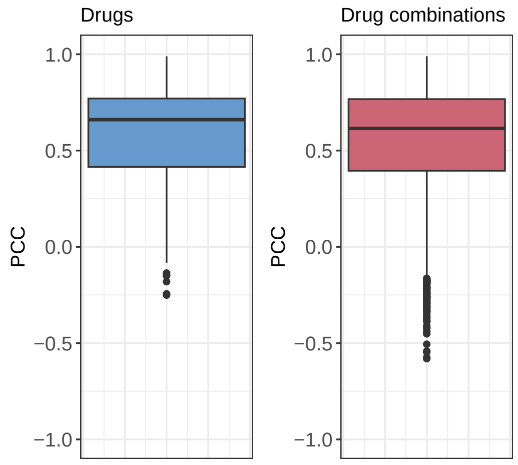


Figure 2: Average Pearson correlation between actual and predicted relative inhibitions per drug (blue) and per drug combination (red).

#### 3 Impact of available Training Data

Since the amount of available data for the investigated drugs and drug combinations differs considerably, we investigated whether the amount of available data has an impact on prediction accuracy. Figure 3A in this Supplement shows a slight negative correlation (PCC -0.24) between the number of available training data points (mono- and combination data) for each drug and its average monotherapy prediction error using the MACCS random forest model. Prediction errors between drugs vary notably, with the largest errors being obtained for drugs with little data. However, also drugs with little training data can achieve small errors.

Interestingly, for drug combinations, the test MAE depends even less on the amount of available combination data (PCC -0.025) or data of the individual drugs (PCC -0.11), as shown in Figures 3B and 3C. This is generally desirable since it shows that the training is not driven only by drugs/combinations with large amounts of data. However, Figure 3 still highlights that certain drugs/combinations can be predicted much more accurately than others.

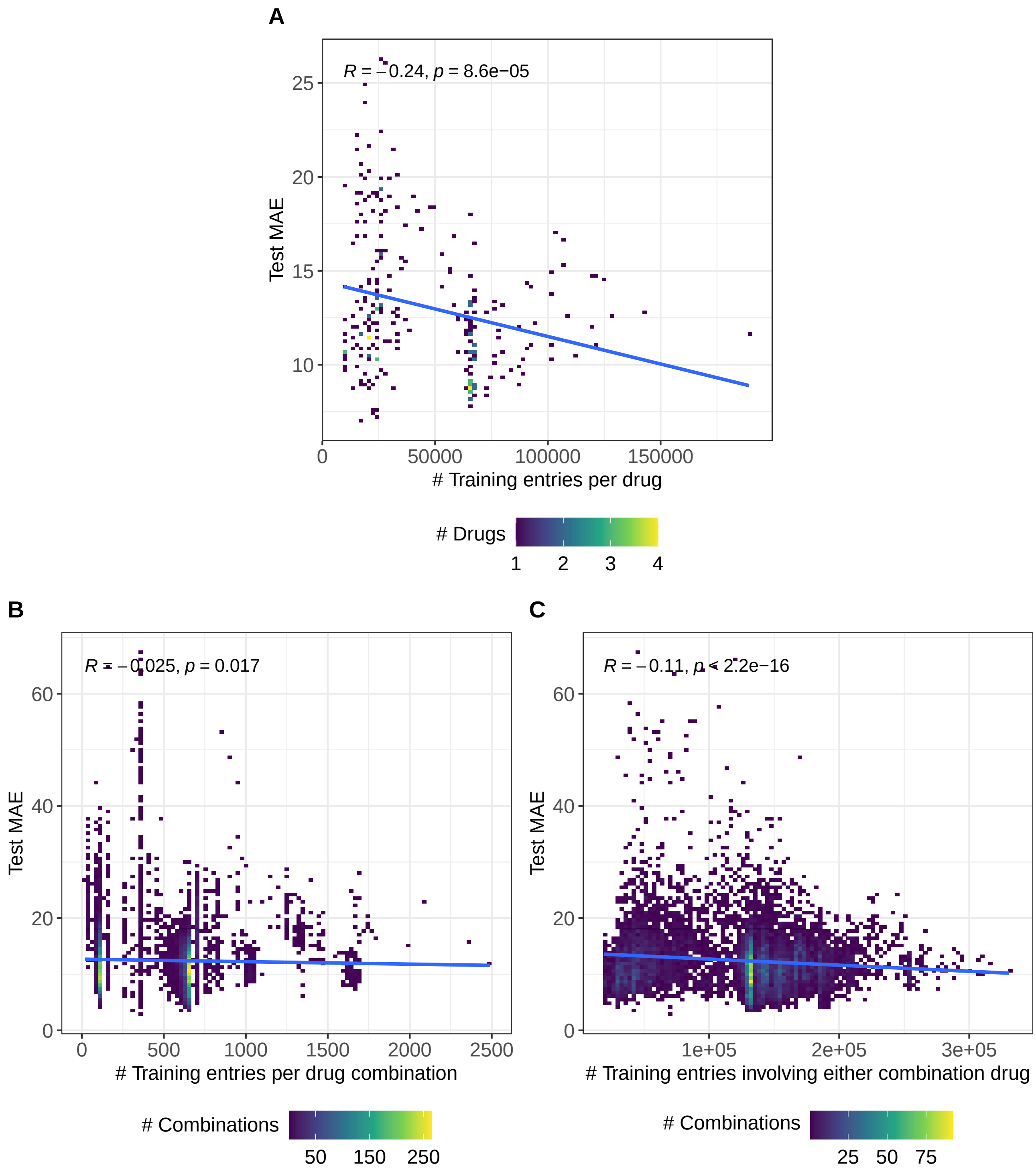


Figure 3: Impact of amount of training entries on predictions. This figure depicts the correlation between the amount of available training data and prediction errors. Sub-figure A shows the correlation between the number of training entries involving a certain drug and its test MAE for monotherapies. Sub-figure B shows the correlation between the number of training entries involving a certain drug combination and its test MAE for combination therapies. Sub-figure C shows the correlation between the number of training entries involving either drug of the investigated combination and its test MAE for combination therapies. In each figure, a linear regression line is shown in blue and R denotes the Pearson correlation.

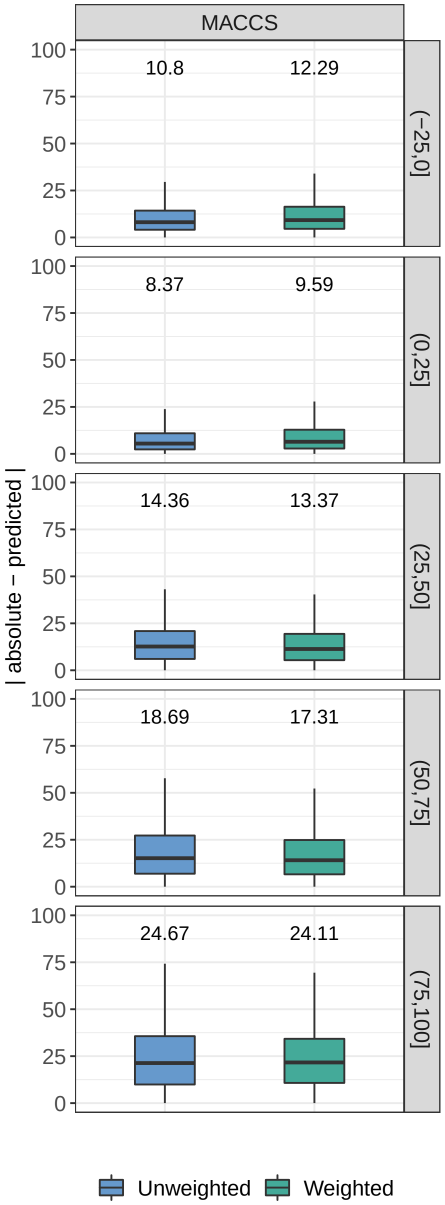


Figure 4: Impact of sample weights on model predictions. This figure compares the prediction performance (in terms of absolute difference between actual and predicted values) for the random forest MACCS model with (green) and without (blue) sample weights. Each row shows the performance for a different interval of actual relative inhibitions. On top of each boxplot, the mean absolute error (MAE) is shown. The weight of each sample was determined based on its interval $i \in I = \{(-\infty, 0], (0, 25], (25, 50], (50, 75], (75, \infty)\}$ as $(\frac{max_{j \in I}|j|}{|i|})^2$, where $|i|$ denotes the number of training samples in interval $i$.

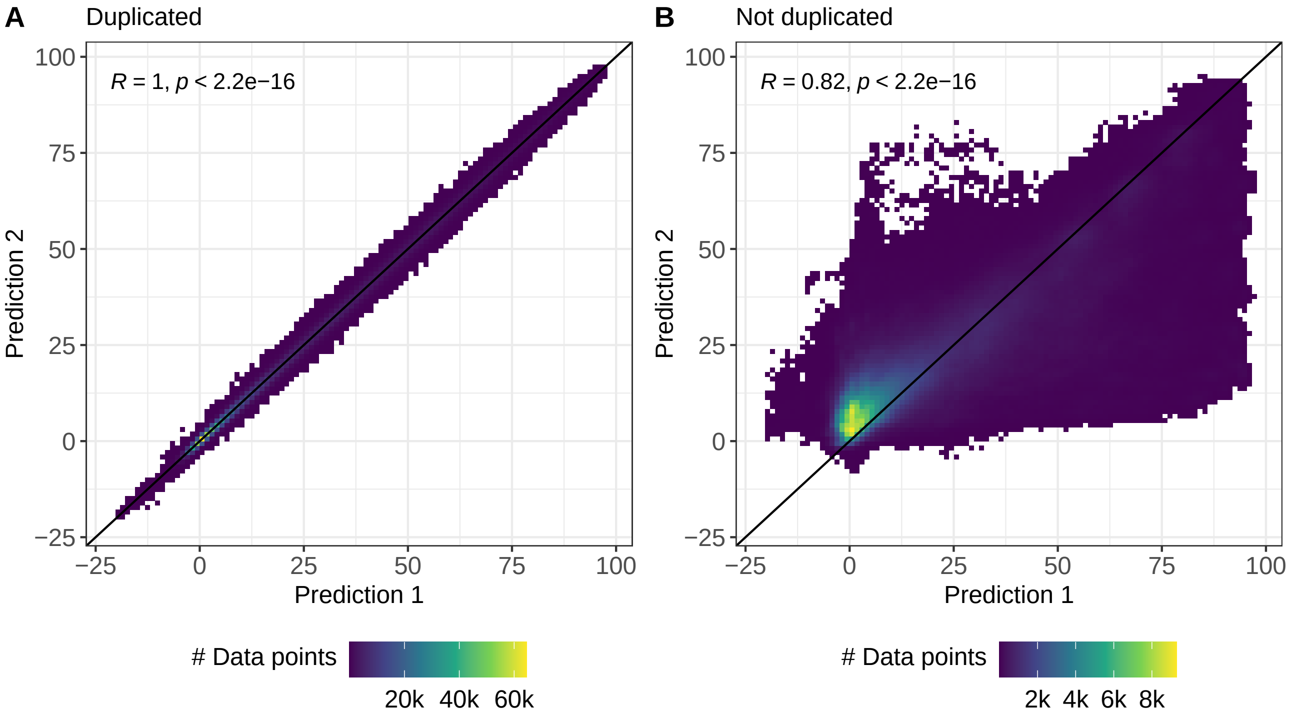


Figure 5: Correlation of duplicated entries from the test data. This figure shows the correlation between the predictions for duplicated entries obtained from the random forest PhysChem model. Duplicated entries refer to the same drug-drug-cell combination and the same treatment concentrations but can be represented by two different model inputs through swapping the features of the respective drugs (cf. Methods and Figure 1 in the main manuscript). Sub-figure A shows the test predictions when including duplicated entries into the training data, while Sub-figure B shows the predictions when training only on non-duplicated entries. In both figures, the black diagonal line represents the identity and R denotes the Pearson correlation between the predictions.

### 4 Additional Analyses Concerning the Reconstruction of Sensitivity Measures

**Hypothesis 1:** The CMax viability is difficult to predict since concentrations exceeding the CMax concentration were not screened for 14 of 77 drugs, corresponding to 30% of the drug-cell line combinations in the test set. Thus, the curve-fitting for these entries might not accurately model the CMax viability.

**Evaluation 1:** We evaluated the PCC only on those drug-cell line combinations for which a concentration greater than CMax was screened. This increased the average PCC slightly from 0.1 to 0.12.

**Hypothesis 2:** Larger prediction errors for high inhibitions (i.e., low viabilities) make the curve-fitting unreliable in areas of high inhibition (cf. Figure 4 of main manuscript), which affects the derived response measures.

**Evaluation 2:** The IC50 value is designed to measure the drug response at a relative inhibition of 50%. To assess the performance at smaller inhibitions (i.e., higher viabilities), we reconstructed IC75 and IC90 values from the fitted curves. The IC75 (IC90) measures the drug concentration where a relative inhibition of $100\% - 75\% = 25\%$ ($100\% - 90\% = 10\%$) is reached. The average per-drug PCCs for the IC75 and IC90 reconstruction are 0.11 and 0.1, respectively, thus, there is no (strong) improvement compared to the IC50 predictions (cf. Figure 6 of this Supplement).

**Hypothesis 3:** The curve-fitting itself might not be beneficial to reconstruct CMax viabilities.

**Evaluation 3:** Instead of deriving the CMax viability from the estimated dose-response curves, we used our model directly to predict the relative inhibition at the CMax concentration and converted this prediction into a relative viability. However, this did also not improve correlations (PCC 0.04, cf. Supplementary Figure 7). Instead, the curve fitting seems to enhance predictions slightly, which is in line with the findings by Rahman and Pal [14].

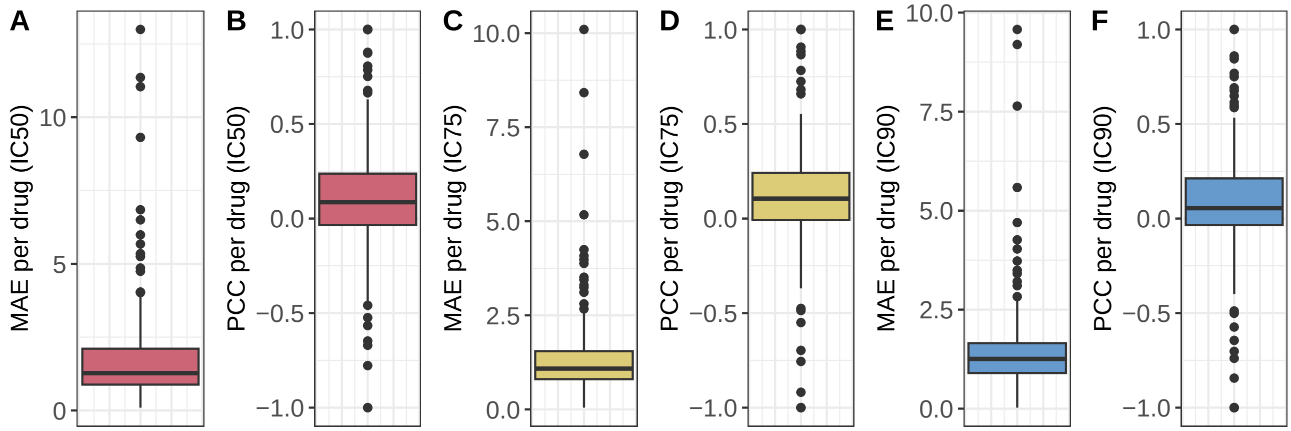


Figure 6: Reconstruction of IC50, IC75, and IC90 values from model predictions. Sub-figures A and B (red) show the distribution of MAE and PCC per drug for the reconstruction of IC50 values using the test set monotherapy data. Sub-figures C and D (yellow) show the analogous results for IC75 values. Sub-figures E and F (blue) show the analogous results for IC90 values.

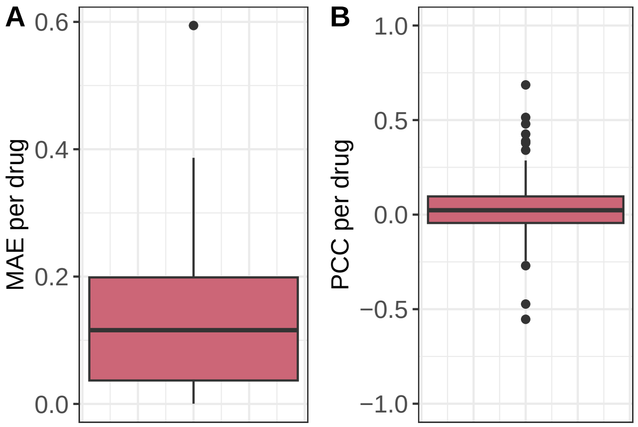


Figure 7: Direct prediction of CMax viabilities. Sub-figures A and B show the distribution of MAE and PCC per drug for the prediction of CMax viabilities using the cell line-drug combinations from the test set monotherapy data.

Table 3: Drug sensitivity/synergy prediction literature. This table lists 39 approaches for drug sensitivity or synergy prediction. For each approach, it is denoted whether drug-specific correlations are provided (Pearson correlation for regression, Matthews correlation for classification) , whether a cell-blind evaluation was performed, whether multi-drug models were trained, and whether the approach explicitly employs dose-response curves (as opposed to mere sensitivity measures like the IC50). Rows marked in light blue denote manuscripts where only classification but no regression was performed. Table continues on next page.

| Model | Correlation per drug | cell-blind | multi-drug | curve-based |
|---|---|---|---|---|
| Menden et al. (2013) [15] | ✗ | ✗ | ✓ | ✗ |
| Zhang et al. (2015) [16] | ✓ | ✗ | ✓ | ✗ |
| Rahman and Pal (2016) [14] | ✓ | ✓ | ✗ | ✓ |
| SRMF (2017) [17] | ✓ | ✗ | ✓ | ✗ |
| HARF (2017) [18] | ✗ | ✓ | ✗ | ✗ |
| TreeCombo (2018) [19] | ✗ | ✗ | ✗ | ✗ |
| RWEN (2018) [20] | ✗ | ✓ | ✗ | ✗ |
| CDRscan (2018) [21] | ✓ | ✗ | ✗ | ✗ |
| QRF (2018) [22] | ✓ | ✓ | ✗ | ✗ |
| NCFGER (2018) [23] | ✓ | ✗ | ✗ | ✗ |
| DeepDR (2019) [24] | ✗ | ✓ | ✗ | ✗ |
| netBITE (2019) [25] | ✓ | ✓ | ✗ | ✗ |
| FRF (2019) [26] | ✗ | ✓ | ✗ | ✓ |
| Sidorov et al. (2019)[27] | ✗ | ✗ | ( ✓) cell line-specific models | ✗ |
| Deng et al. (2020) [28] | ✗ | ( ✓) LOOCV / train on CCLE, test on GDSC | ✗ | ✗ |
| Ahmed et al. (2020) [29] | ✓ | ✓ | ✗ | ✗ |
| ADRML (2020) [30] | ✓ | ✗ | ✓ | ✗ |
| Julkunen (2020) [31] | ✗ | ✗ | ✓ | ✗ |
| MinDrug [32] | ✗ | ✓ | ( ✓) subsets of similar drugs | ✗ |
| PathDSP (2021) [33] | ( ✓) not cell-blind | ✓ | ✓ | ✗ |
| REFINED CNN (2021) [34] | ✗ | ✗ | ( ✓) cell line-specific models | ✗ |
| GraphDRP (2021) [35] | ✗ | ✓ | ✓ | ✗ |
| Precily (2022) [36] | ✓ | ✓ | ✓ | ✗ |
| KBMTL (2014) [37] | ✗ | ✓ | ✓ | ✗ |
| DeepSynergy (2018) [38] | ✓ | ✗ | ✓ | ✗ |
| DeepCDR (2020) [39] | ( ✓) not cell-blind | ✓ | ✓ | ✗ |
| Kim et al. (2021) [40] | ✗ | ✗ | ✓ | ✗ |
| MatchMaker (2022) [41] | (✗) drug pairs | ✗ | ✓ | ✗ |
| SAURON-RF (2022) [42] | (✓) only classification | ✓ | ✗ | ✗ |
| reliable SAURON-RF (2023) [43] | ✓ | ✓ | ✗ | ✗ |
| LOBICO (2016) [44] | ✗ | ✓ | ✗ | ✗ |
| Stanfield et al. (2017) [45] | ✗ | ✗ | ( ✓) train on CCLE, test on GDSC | ✗ |
| SyDRa (2017) [46] | ✗ | ( ✓) train of DREAM, test on CMap | ✓ | ✗ |

Continuation of Table 3

| **Model** | **Correlation per drug** | **cell-blind** | **multi-drug** | **curve-based** |
|---|---|---|---|---|
| SyDRa (2017) [46] | ✗ | (✓) train of DREAM, test on CMap | ✓ | ✗ |
| HNMDRP (2018) [47] | ✗ | ✗ | ✓ | ✗ |
| Deep-Resp-Forest (2019) [48] | ✗ | ✓ | ✗ | ✗ |
| MOLI (2019) [49] | ✗ | (✓) train on GDSC, test on TCGA + PDX | (✓) drugs with same target pathway | ✗ |
| MERIDA (2021) [50] | ✗ | ✓ | ✗ | ✗ |
| RAMP (2022) [51] | ✗ | (✓) patient data | ✓ | ✗ |
| NeRD (2022) [52] | ✗ | ✓ | ✓ | ✗ |
| GADRP (2023) [53] | ✗ | ✗ | ✓ | ✗ |

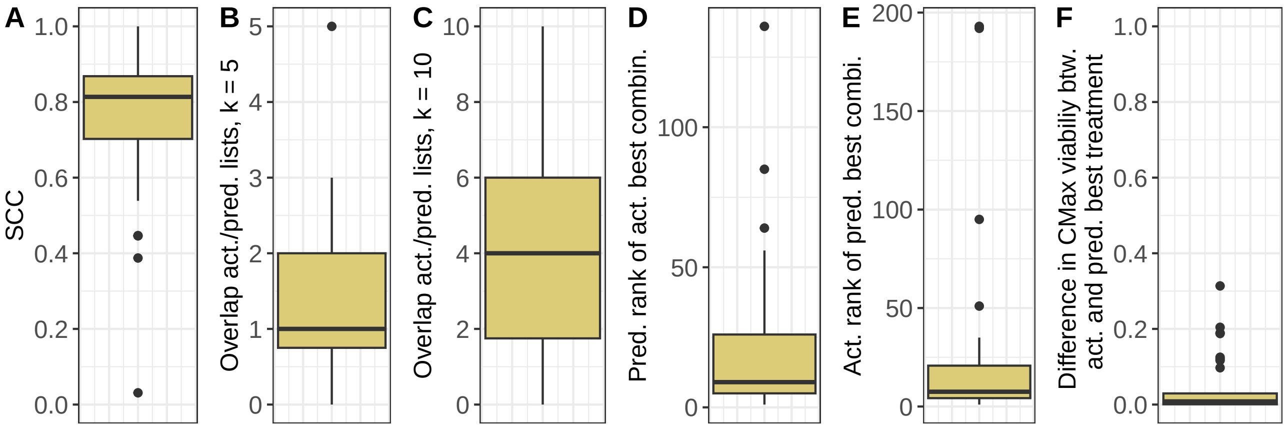


Figure 8: Treatment prioritization. This figure depicts the test set prioritization results for combination therapies including (A) the SCC between the actual and predicted rankings for each cell line, (B)/(C) the intersection size between the 5/10 actual and predicted most effective treatments, (D) the predicted rank of the actual most effective treatment, (E) the actual rank of the treatment predicted to be most effective, and (F) the difference between the actual CMax viabilities for the actual and predicted most effective treatment.

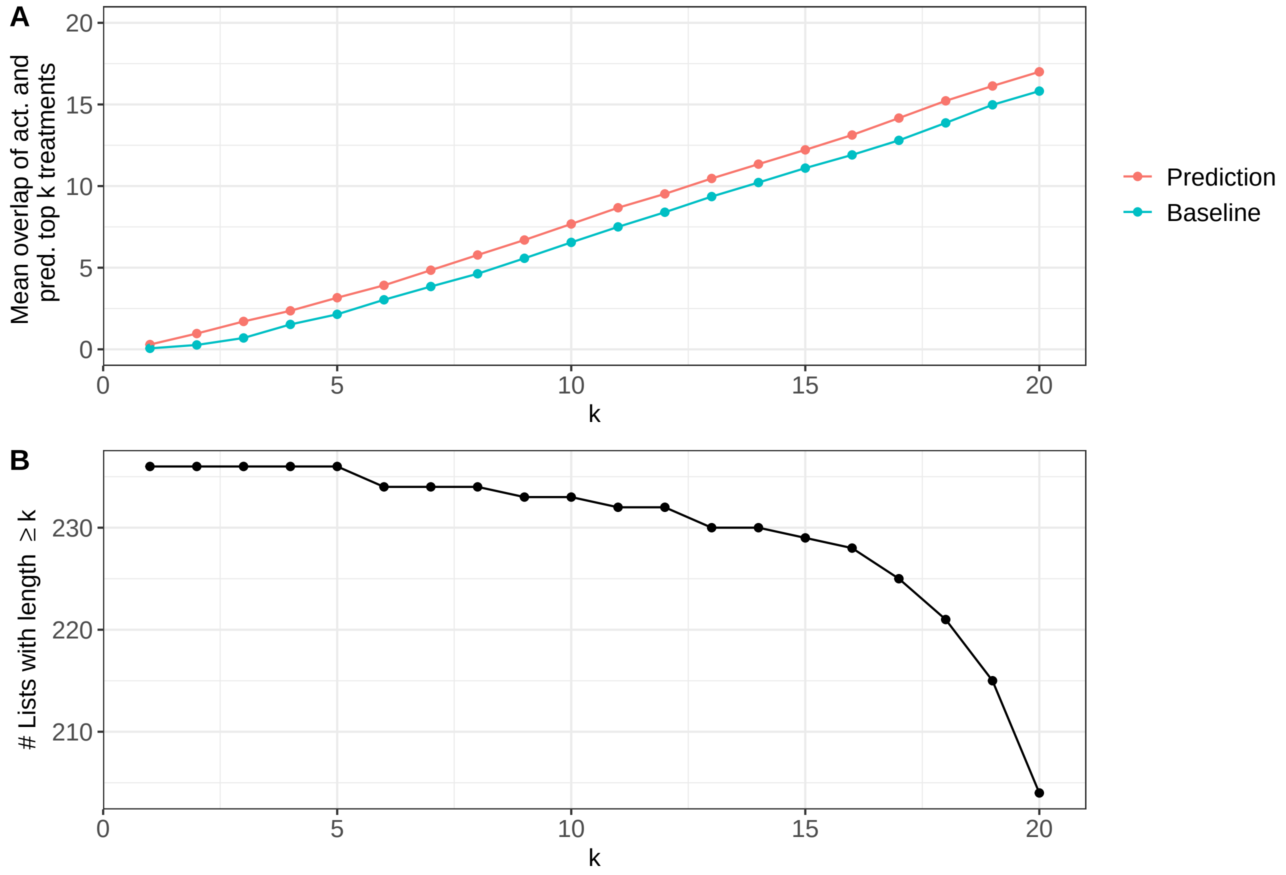


Figure 9: Overlap of $k$ actual and predicted best treatments for monotherapies. Sub-figure A shows the average intersection size between the $k$ actual best treatments and the $k$ predicted best treatments for each cell line. Sub-figure B shows the number of cell lines based on which the average for each $k$ was computed.

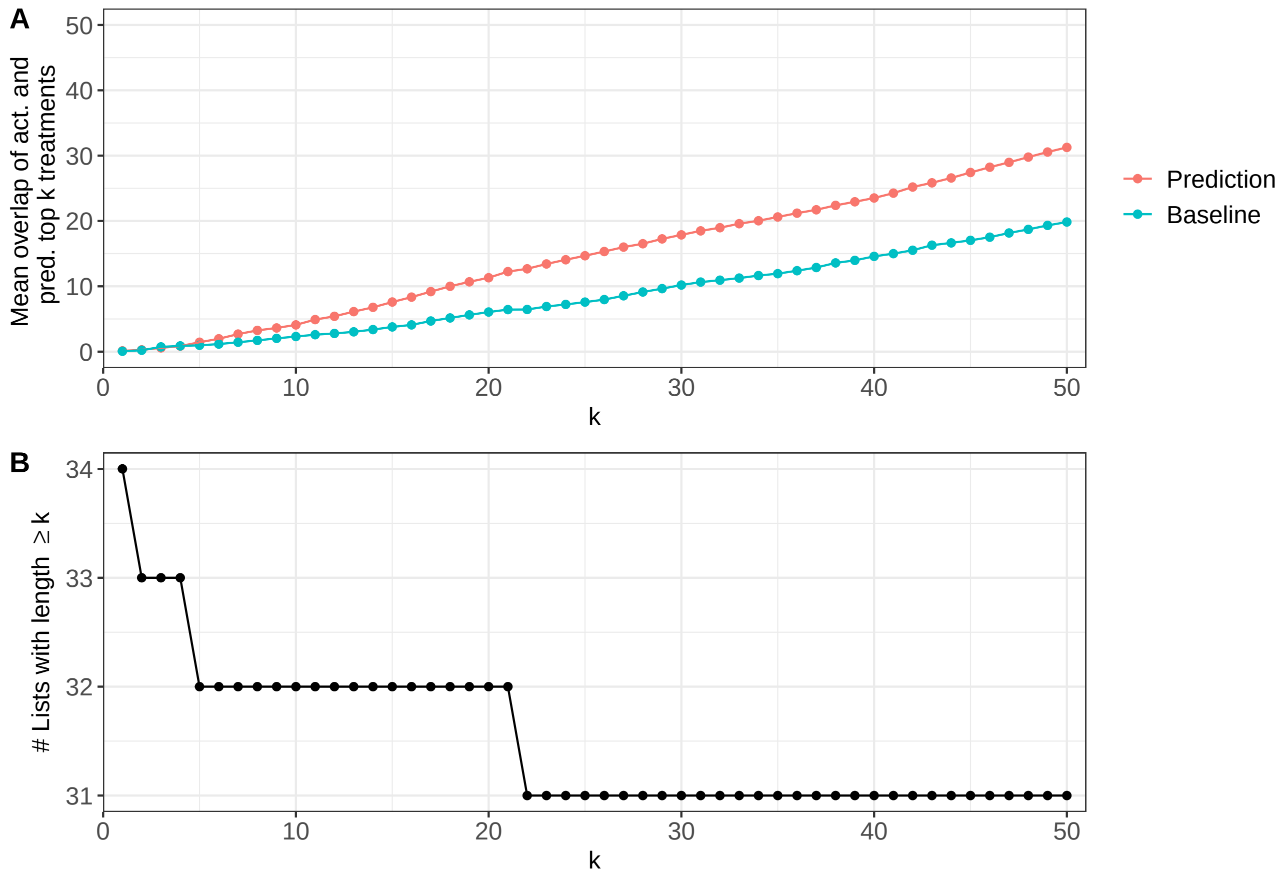


Figure 10: Overlap of $k$ actual and predicted best treatments for combination therapies. Sub-figure A shows the average intersection size between the $k$ actual best treatments and the $k$ predicted best treatments for each cell line. Sub-figure B shows the number of cell lines based on which the average for each $k$ was computed. Note that the number of test cell lines with available combination data is smaller than the number of cell lines with available monotherapy data.

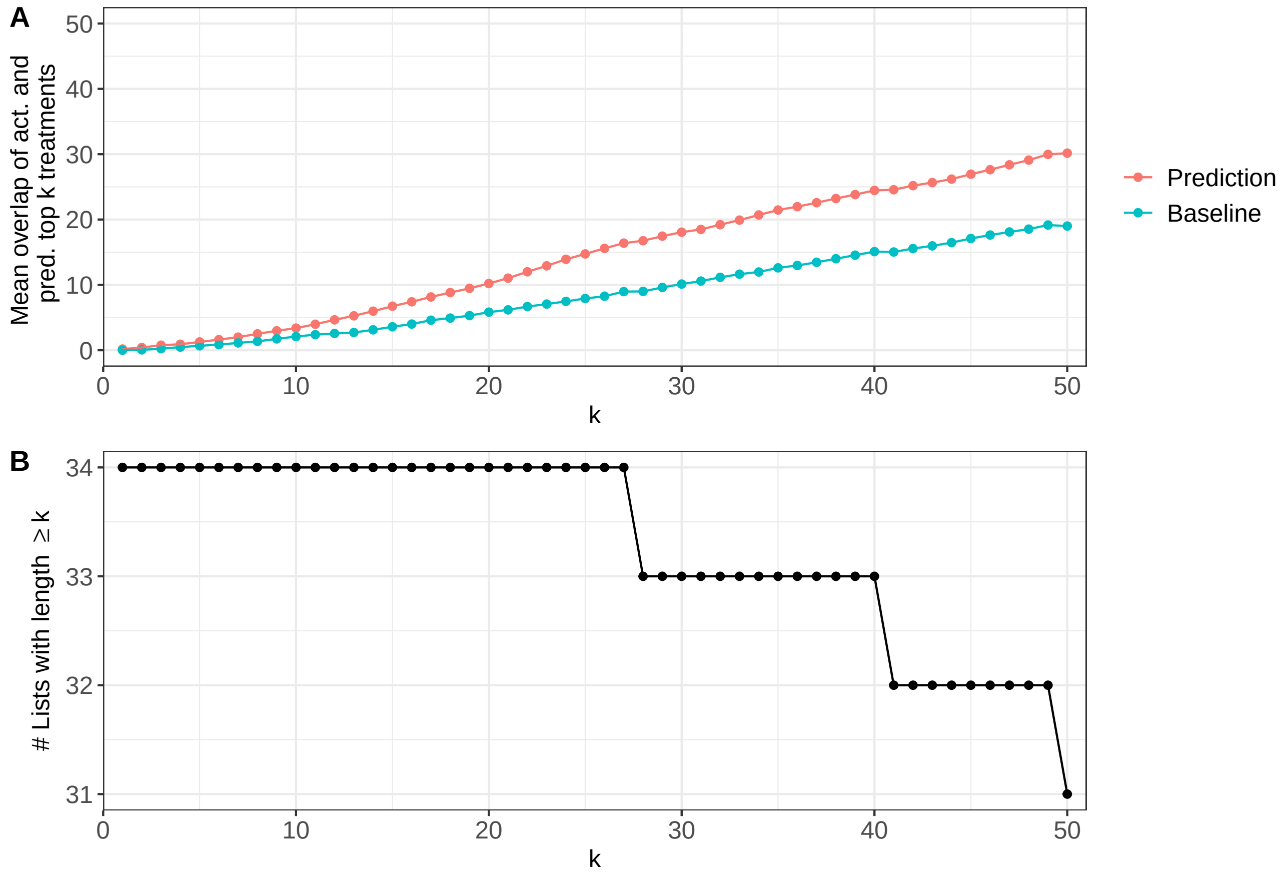


Figure 11: Overlap of $k$ actual and predicted best treatments for the combination of both mono- and combination therapies. Sub-figure A shows the average intersection size between the $k$ actual best treatments and the $k$ predicted best treatments for each cell line. Sub-figure B shows the number of cell lines based on which the average for each $k$ was computed. Note that the number of test cell lines with available combination data is smaller than the number of cell lines with available monotherapy data and we show the results only for cell lines where both were available.

[11] Shuyu Zheng, Wenyu Wang, Jehad Aldahdooh, Alina Malyutina, Tolou Shadbahr, Alberto Pessia, and Jing Tang. Synergyfinder plus: towards a better interpretation and annotation of drug combination screening datasets. *bioRxiv*, page 2021.06.01.446564, 2021.

[19] Joseph D Janizek, Safiye Celik, and Su-In Lee. Explainable machine learning prediction of synergistic drug combinations for precision cancer medicine. *BioRxiv*, page 331769, 2018.

[34] Omid Bazgir, Souparno Ghosh, and Ranadip Pal. Investigation of REFINED CNN ensemble learning for anti-cancer drug sensitivity prediction. *Bioinformatics*, 37(Supplement 1):i42–i50, 07 2021.

[35] Tuan Nguyen, Giang TT Nguyen, Thin Nguyen, and Duc-Hau Le. Graph convolutional networks for drug response prediction. *IEEE/ACM transactions on computational biology and bioinformatics*, 19(1):146–154, 2021.

[36] Smriti Chawla, Anja Rockstroh, Melanie Lehman, Ellca Ratther, Atishay Jain, Anuneet Anand, Apoorva Gupta, Namrata Bhattacharya, Sarita Poonia, Priyadarshini Rai, et al. Gene expression based inference of cancer drug sensitivity. *Nature communications*, 13(1):5680, 2022.

[37] Mehmet Gönen and Adam A Margolin. Drug susceptibility prediction against a panel of drugs using kernelized bayesian multitask learning. *Bioinformatics*, 30(17):i556–i563, 2014.

[38] Kristina Preuer, Richard PI Lewis, Sepp Hochreiter, Andreas Bender, Krishna C Bulusu, and Günter Klambauer. Deepsynergy: predicting anti-cancer drug synergy with deep learning. *Bioinformatics*, 34(9):1538–1546, 2018.

[39] Qiao Liu, Zhiqiang Hu, Rui Jiang, and Mu Zhou. Deepcdr: a hybrid graph convolutional network for predicting cancer drug response. *Bioinformatics*, 36(Supplement_2):i911–i918, 2020.
